# SQANTI3: curation of long-read transcriptomes for accurate identification of known and novel isoforms

**DOI:** 10.1101/2023.05.17.541248

**Authors:** Francisco J. Pardo-Palacios, Angeles Arzalluz-Luque, Liudmyla Kondratova, Pedro Salguero, Jorge Mestre-Tomás, Rocío Amorín, Eva Estevan-Morió, Tianyuan Liu, Adalena Nanni, Lauren McIntyre, Elizabeth Tseng, Ana Conesa

## Abstract

The emergence of long-read RNA sequencing (lrRNA-seq) has provided an unprecedented opportunity to analyze transcriptomes at isoform resolution. However, the technology is not free from biases, and transcript models inferred from these data require quality control and curation. In this study, we introduce SQANTI3, a tool specifically designed to perform quality analysis on transcriptomes constructed using lrRNA-seq data. SQANTI3 provides an extensive naming framework to describe transcript model diversity in comparison to the reference transcriptome. Additionally, the tool incorporates a wide range of metrics to characterize various structural properties of transcript models, such as transcription start and end sites, splice junctions, and other structural features. These metrics can be utilized to filter out potential artifacts. Moreover, SQANTI3 includes a Rescue module that prevents the loss of known genes and transcripts exhibiting evidence of expression but displaying low-quality features. Lastly, SQANTI3 incorporates IsoAnnotLite, which enables functional annotation at the isoform level and facilitates functional iso-transcriptomics analyses. We demonstrate the versatility of SQANTI3 in analyzing different data types, isoform reconstruction pipelines, and sequencing platforms, and how it provides novel biological insights into isoform biology. The SQANTI3 software is available at https://github.com/ConesaLab/SQANTI3.

## **1** Introduction

Long-read sequencing has been declared the Method of the Year 2022 by Nature Methods [1]. The ability of technologies like Pacific Biosciences (PacBio) and Oxford Nanopore Technologies (ONT) to obtain kilobase-range reads from single molecules has opened up new research opportunities in the study of genomes and transcriptomes. When applied to the analysis of gene expression, long-read RNA sequencing (lrRNA-seq) has the potential to capture full-length transcripts and reveal isoform diversity. This approach has been used to profile complex transcriptomes from neural tissues [2], tumor cells [3], T-cell receptors [4], and multiple plant tissues [5]. In addition, lrRNA-seq has been used to study allele-specific gene expression [6], circular RNAs [7], functional pseudogenes [8], and single-cell isoform diversity [9] [4].

Motivated by these advances, lrRNA-seq is increasingly being used to study transcriptomes of organisms with poorly annotated transcriptomes and without reference genomes [10], [11], [12]. To support this trend, new software tools have been developed to analyze lrRNA-seq data, including methods for transcript identification [13], [14] [15], quantification [16] [17], differential splicing analysis [18], and functional interpretation of long-read data [19].

One of the most striking results of lrRNA-seq utilization is the identification of thousands of novel transcripts, even in well-annotated genomes [16]. However, long-read data is not error-free, and some of these transcript models may represent false calls. While the initial high levels of noise of both PacBio and ONT,[20] have been reduced, respectively, by the introduction of HiFi reads and improved base calling[21] [22], other types of errors still exist in lrRNA-seq datasets, such as those caused by RNA degradation, library preparation, and read mapping. Adequate data quality control is critical to realize the full potential of these technologies.

SQANTI (Structural and Quality Annotation of Novel Transcript Isoforms) was published in 2018 as a tool for quality control of long-read transcriptomics datasets. The software compared long-read-derived transcript models to a reference annotation and classified them as Full Splice Match (FSM), Incomplete Splice Match (ISM), Novel In Catalogue (NIC), Novel Not in Catalogue (NCC), Fusion, Antisense, Intronic, Genic, Genomic, or Intergenic [23] (Fig.1**a**). SQANTI also extracted over 50 features that described the characteristics of the predicted transcript models and uses them to eliminate potential artifacts through a Machine Learning approach. SQANTI was quickly adopted by the lrRNA-seq community, and its structural classification of isoforms has become standard for describing the composition of long-read-defined transcriptomes [24] [25] [26] [27].

**Fig. 1.**
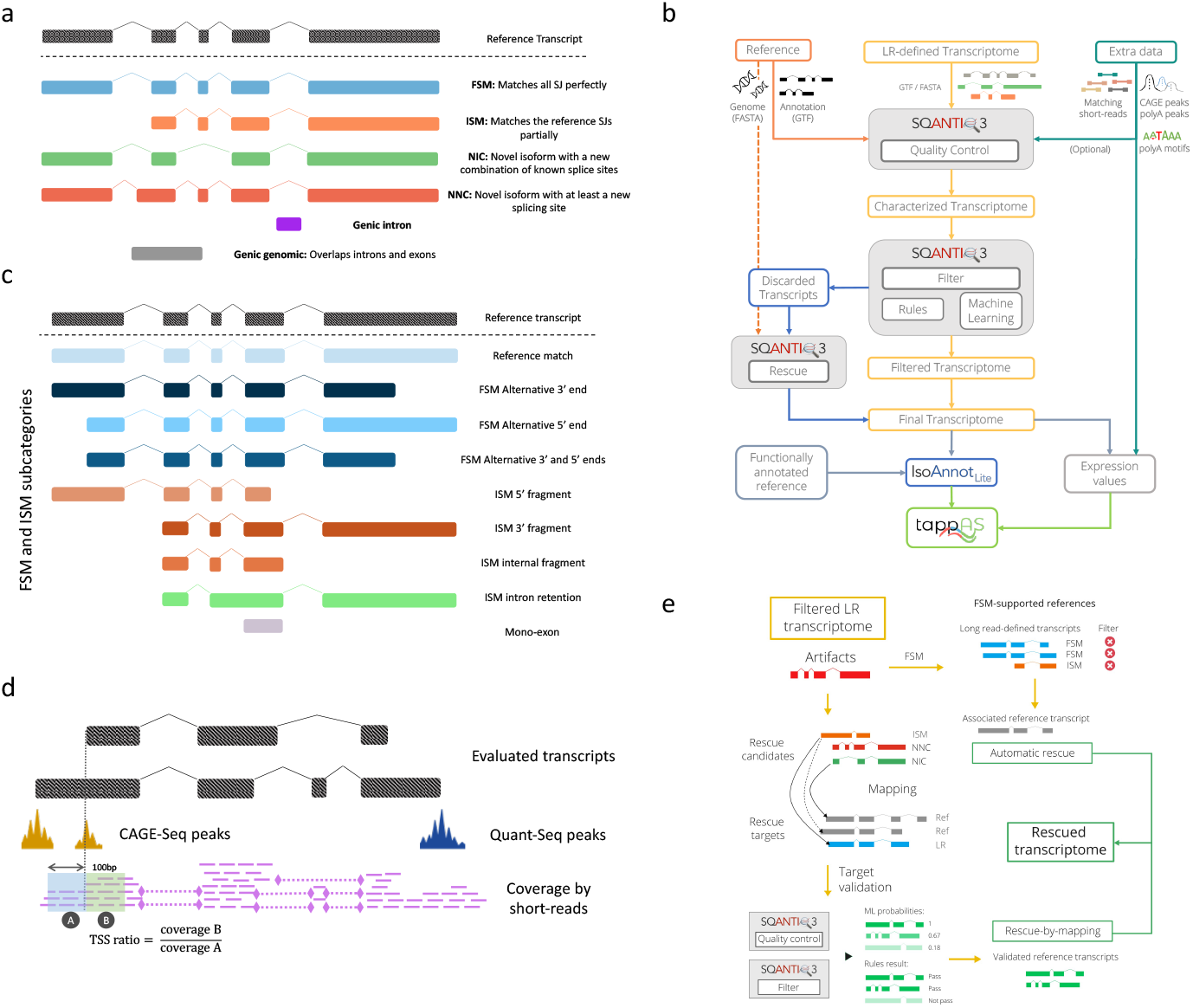
Overview of SQANTI3. **a** Main SQANTI structural categories for transcript models of known genes, which are based only on the completeness and novelty status of the string of SJ detected. **b** SQANTI3 workflow is divided into three steps: Quality Control, Filter, and Rescue. The final transcriptome can be functionally annotated with IsoAnnotLite for downstream analyses. **c**, SQANTI3 subcategories for FSM and ISM isoforms. **d** Orthogonal data features. **e**Schematic representation of the novel Rescue function.

The initial SQANTI quality control (QC) and filtering modules were designed to identify low-quality transcripts models based on the characteristics of their splice junctions, as high error rates in long-read technologies were expected to result in false splice sites. However, the field’s understanding of lrRNA-seq data limitations has grown with the usage of the technology in studies with increasing complexity and scale. For example, lrRNA-seq transcriptomes have been found to exhibit important variability at the 3’ and 5’ ends, which may reflect alternative transcription start or terminating sites (TSS and TTS, respectively) or be a consequence of library artifacts and RNA degradation. Moreover, additional data such as CAGE [28], Quant-seq [29], and Illumina reads [30] might be available and can be used to further characterize, validate and eventually correct long read transcript models. Hence, a state-of-the-art quality control strategy for lrRNA-seq should capture these aspects of transcript diversity, incorporate multiple sources of information in the characterization of the transcript models and effectively use this information to improve the quality of the transcriptomes inferred by long-read sequencing.

Here we present SQANTI3 (SQ3), an open-source and collaborative project for the extensive assessment, curation, and annotation of lrRNA-seq transcriptomes. SQANTI3 extends the original transcript classification scheme to better reflect 3’ and 5’ end diversity and incorporates multiple data sources to expand the number of QC metrics calculated by the tool. Additionally, SQANTI3 contains improved strategies for artifact removal and a novel Rescue functionality that suggests a representative isoform to replace discarded transcript models, thereby improving the accuracy and completeness of longread-defined transcriptomes. SQANTI3 also integrates IsoAnnotLite, a tool for the functional annotation of isoforms, which facilitates downstream analyses of transcriptome biology. The SQANTI3 toolkit is technology and analysis pipeline-independent, capable of characterizing data from both PacBio and ONT platforms. The method has been used in numerous lrRNA-seq studies and is the evaluation engine of the LRGASP challenge, an international benchmarking initiative where over 130 long-read transcriptomes obtained from a wide range of experimental and bioinformatics methods were assessed and compared [31]. The expansion of capabilities in SQANTI3 reflects the community’s growing need for advanced bioinformatics tools for the evaluation and curation of transcriptomes.

## 2 Results

### 2.1 SQANTI3 workflow overview

SQANTI3 enhances the capabilities of SQANTI by incorporating 18 new functionalities, thereby offering a comprehensive toolkit for improving long-read-defined transcriptomes. These additions comprise new structural categories, additional quality descriptors of transcript start and end sites, and two extra tools for lrRNA-seq isoform filtering and transcriptome curation (see Supplementary Table A1 for a comparison between SQANTI and SQANTI3). The SQANTI3 workflow includes three modules (Fig.1**b**). The Quality Control (QC) module computes SQANTI3 descriptors and structural categories to characterize long-read-defined transcriptomes. The Filter module provides two approaches to identifying transcript models that are potential artifacts: a rules filter and a machine learning filter. The Rescue module re-evaluates artifacts to suggest a *bona fide* replacement transcript model and avoid the loss of known genes and transcripts for which evidence of expression exists.

Additionally, SQANTI3 enables the functional annotation of transcript models through the integration of IsoAnnotLite, which maps a pre-computed database of transcripts annotated with functional domains and motifs to the long-read transcript sequences. The result is a curated and functionally annotated fulllength transcriptome in a file format compatible with the tappAS software [19] for the visualization and assessment of isoform expression changes (Fig.1**b**).

#### Novel structural categories to describe variability at the 3’ and 5’ ends

SQANTI3 defines structural sub-categories of Full (FSM) and Incomplete Splice Matches (ISM) to more accurately capture variability at the transcript ends (Fig.1**c**). The Reference Match (RM) subcategory is defined as an FSM where both the 3’ and 5’ ends are within 50bps of the reference transcript’s TTS and TSS, respectively. RMs, therefore, match their associated reference transcripts both at junctions and end sites and can be considered complete hits when comparing a lrRNA-seq transcriptome to the reference. Conversely, non-RM FSM transcripts show 3’ and 5’ end variability within the first or last exon of the reference transcript and are included within the alternative 3’ and/or 5’ end categories. Similarly, SQANTI3 divides ISM transcripts into subcategories based on whether missing splice junctions (SJ) are located at the 5’ (3’ fragment), 3’ (5’ fragment), or both ends (internal fragment). If SJs are lost due to an intron retention event, these are labeled accordingly. Finally, since mono-exonic transcripts cannot be analyzed using SJ-centric criteria, this subcategory remains the same across all structural categories (Fig.1**c**).

#### SQANTI3 validates 3’ and 5’ ends with additional data

To assess the reliability of TSS and TTS of long-read transcript models, SQANTI3 QC can now process complementary data such as CAGE, Quant-seq, or other genomic-region data (Fig.1**d**) and calculate the distance and overlap between the 3’ and 5’ transcript ends and these regions. The software also includes automation of short-read data processing as part of the QC workflow and introduces an additional TSS-related metric, the TSS ratio. Briefly, for each transcript, the ratio between short-read coverage downstream and upstream of the TSS is computed by SQANTI3 QC (see Methods). 5’ end-degraded transcripts are expected to have uniform coverage on both sides of the TSS (i.e., TSS ratio*≈*1), while a true TSS is expected to have much lower upstream coverage, resulting in a TSS ratio greater than one.

#### Novel artifact detection strategies improve versatility in QC

SQANTI3 includes a Random Forest Machine Learning (ML) classifier that labels long-read transcript models as isoforms or artifacts using SQANTI QC descriptors as predictive variables and a set of user-defined true and false transcripts for model training. This machine learning method has been substantially updated to include the new quality features, particularly those derived from additional data, allow more flexibility in the selection of predictor variables and improve overall usability (Supplementary Table A1). Since defining a training set might not always be feasible, SQANTI3 also alternatively offers a Rules filter function. This function allows users to easily apply exclusion criteria based on the QC descriptors, providing flexibility in designing a curation strategy that can be adapted to the available data and the characteristics of the algorithm used to infer transcript models. Moreover, SQANTI3 Filter is now fully integrated within the SQANTI3 curation pipeline and provides extensive reporting of filtering results.

#### Novel rescue module to recover low-quality transcript models

Upon SQANTI3 filtering of long-read transcriptomes known genes may be completely removed when all their transcripts are classified as artifacts. To mitigate the risk of excluding transcripts and genes that show evidence of expression SQANTI3 includes a Rescue module where artifacts are assigned to the most suitable reference or long-read transcript applying a two-step process (Fig.1**e**, Supplementary Fig.A1). Reference transcripts with FSM-based evidence for which all associated isoforms have been removed are recovered, i.e. *automatic rescue*. ISM, NIC, and NNC artifacts (rescue candidates) are compared to both reference and long-read defined transcript models (rescue targets) to identify, where possible, matching transcripts. These rescue targets are evaluated using the same selection criteria applied during the filtering step and, if not already present, the best target transcript model is added to the reference transcriptome.

### 2.2 SQANTI3 QC of a WTC11 human cell-line dataset

To demonstrate and validate SQANTI3, we used cDNA-PacBio data from the human WTC11 cell line obtained by the LRGASP project [31]. Illumina, CAGE-seq, and Quant-seq data were also available for this cell line. Transcript models were obtained from the long reads using IsoSeq3 (see Methods), which is a reference-independent method widely used by the PacBio transcriptomics community that did not participate in LRGASP. A total of 228,379 transcript models were built, of which 209,220 belonged to 17,467 known genes (Fig.2**a**). Almost one-third of all transcripts were classified as ISM isoforms (67,804; 29.69%), while 37% of them were described as novel isoforms of annotated genes (48,878 NIC and 35,743 NNC). Only 56,795 transcripts (24.87%) were annotated under the FSM category, a small fraction considering that WTC11 is a well-studied human cell line. While this may be due to reference catalog incompleteness, it could also be attributed to inaccurately defined isoforms resulting from errors in the long-read technology or transcript reconstruction algorithm. This highly diverse transcriptome dataset provides an excellent testbed for evaluating the effectiveness of SQANTI3 QC in characterizing isoform structural variability and identifying potential sources of artifacts.

**Fig. 2.**
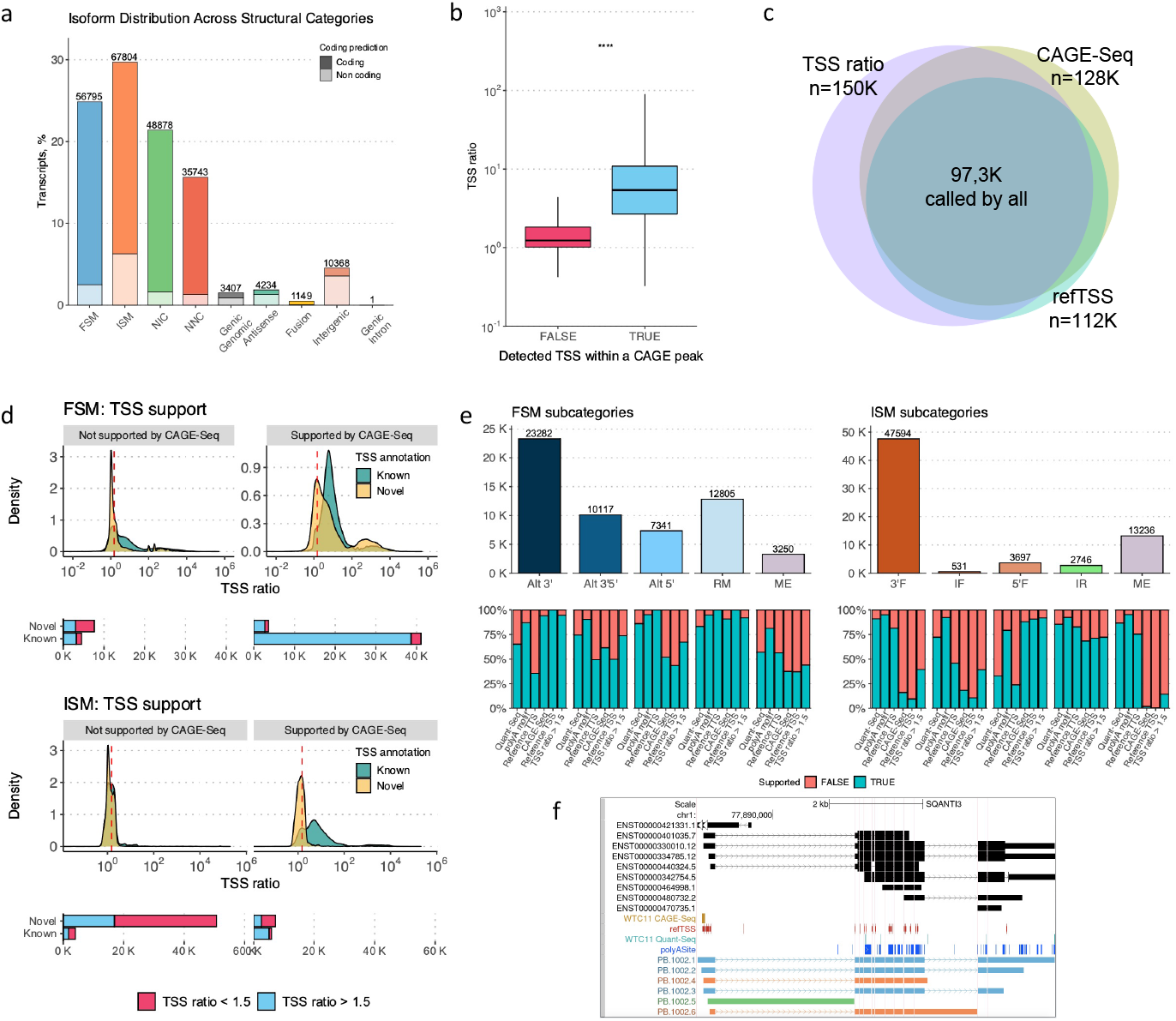
Validation of novel features in SQANTI3. **a** Distribution of transcript models across SQANTI3 structural categories in the WTC11 IsoSeq3-defined transcriptome. **b** TSS ratio difference between TSSs supported and not supported by CAGE-seq data. **c** Agreement in TSS validation using various sources of additional information: TSS ratio based on matching short-reads RNA-seq, sample-specific CAGE-seq, and the refTSS database. **d** Uneven support of CAGE-Seq data for FSM and ISM transcript models, with supported TSSs usually having a TSS ratio higher than 1.5 (red dashed line), particularly if they are also known TSSs. **e** Supporting evidence for FSM and ISM subcategories at 3’ and 5’ ends, respectively, by Quant-seq, presence of polyA motifs, and reference annotation for the former, and CAGE-seq, reference annotation, and TSS ratio for the latter. **f** UCSC Browser view of IsoSeq3 transcripts in the NEXN gene, cataloged as FSM (blue), ISM (orange), and NIC (green) by intron retention. The upper track shows the GENCODE annotation for NEXN (black), while the middle tracks represent CAGE-Seq (gold), refTSS (dark red), Quant-seq (light blue), and polyA site (dark blue) annotations.

Previous versions of SQANTI were targeted to the QC of NIC and NNC isoforms, where differences exist with respect to the reference splice junctions. In this section, we demonstrate how SQANTI3 adds functionality in evaluating the 3’ and 5’ ends of lrRNA-seq transcript models, which is relevant for the WTC11 dataset given the high proportion of ISM in the IsoSeq3 output. We first assessed the reliability of the TSS ratio metric for accurately identifying transcription start sites (TSSs) by comparing the ratio values calculated by SQANTI3 with empirical evidence obtained from CAGE-seq. Among all the identified TSSs, those with additional support from sample-specific CAGE data showed a significantly higher TSS ratio (p-value = 2e-16, Wilcoxon Test, Fig.2**b**). Notably, a TSS ratio threshold *>*1.5 was sufficient to confirm the start sites of 88.2% of transcripts supported by CAGE-seq. Additionally, annotations in the refTSS database [32](i.e., not sample-specific) supported 92.6% of the TSSs with ratios over this threshold (Fig.2**c**). Ultimately, TSS sites with a ratio higher than 1.5 were significantly more likely to be supported by additional data than lower ratio sites (p-value *<* 2e-16, Fisher’s Exact test; see Methods). These results indicate that the proposed TSS ratio metric is a reasonable short-reads-based alternative to CAGE-seq support.

To understand the relationship between TSS-related attributes and transcript length completeness, we compared TSS metrics for transcript models in the FSM and ISM categories. While the vast majority of FSM transcripts showed both CAGE-seq support and TSS ratios larger than 1.5 (Fig.2**d**, top panel), the ISM category was enriched in transcripts without an overlapping CAGE-seq peak (Fig.2**d**, bottom panel). We identified 7,599 ISM isoforms with a reliable TSS according to both CAGE-seq data and TSS ratio, one-third of which were novel TSS with respect to the reference annotation. Conversely, 4,591 FSM isoforms showed low TSS ratios and lacked support from the reference annotation or CAGE-seq data, suggesting that the TSS of these transcript models might not be correctly defined.

For the 3’ end of transcripts, SQANTI3 QC calculates three quality control metrics: the distance to the closest same-gene annotated TTS, the presence of a polyadenylation (polyA) motif within the transcript sequence, and, if available, the support from Quant-seq data. We analyzed the relationship among these metrics. The detection of a polyA motif was found to be significantly associated with 3’ end validation by Quant-seq (p-value = 2.2 e-16, Fisher’s Exact Test, see Methods and Supplementary Fig.A2a). Furthermore, the detected motifs were concentrated at a distance of 16-18 base pairs (bp) from the end of the transcript (as expected based on available experimental evidence [33]) even in those transcripts that were not validated with Quant-seq (Supplementary Fig.A2b). This suggests that long-read sequencing methods detect alternative 3’ ends with higher sensitivity than Quant-seq, possibly because Quant-seq requires high expression of the alternative TTS isoform to call a polyA site. Transcripts that were likely to be a product of intrapriming (defined as 60% As downstream of the TTS [23]), rarely contained a polyA motif or the polyA was located closer to the 3’end than expected (Supplementary Fig.A2c). Overall, these results further support polyA motif detection as a reliable indicator of a *bona fide* transcription termination site.

The results above demonstrate the usefulness of SQANTI3 QC for evaluating the 3’ and 5’ ends of lrRNA-seq transcript models. However, they also suggest that conducting a more in-depth analysis of TSS/TTS variability patterns is advisable, which can be achieved through the novel FSM and ISM subcategories. Interestingly, only 22.5% (12,805 out of 56,795) of FSM were Reference Match (RM) transcripts, while 58.8% of them showed variation at 3énds (belonged to either the Alternative 3’ end or Alternative 5’/3énd subcategories) (Fig. 2**e**). Complementary evidence indicated that these end sites were validated in 39.7%, 67.9%, and 87.8% of cases by a same-gene annotated TTS, Quant-seq data, and the presence of a polyA motif, respectively. A similar pattern was observed for TSS within the Alternative 5’ end subcategories: 47.2% fell within an alternative annotated start site, 66.2% were supported by CAGE-seq, and 70.9% had a TSS ratio greater than 1.5 (Fig.2**e**). The subcategory-level analysis of FSM, therefore, reveals incompleteness in reference annotation and suggests that novel combinations of known start/end sites and intron chains are yet to be described.

The analysis of ISM subcategories showed that 3’ Fragment (3’F) was by far the most abundant group with a total of 47,594 transcript models (70.2% of ISM), where only 9.4% had a known TSS, 16% had CAGE-seq support and 39.5% displayed an above-threshold TSS ratio (Fig.2**e**). This pattern was recapitulated by the 13,236 mono-exonic ISM transcripts, for which most 3’ ends were validated by orthogonal data, whereas 5’ ends remained largely unsupported (Fig.2e). Moreover, transcripts from the mono-exon subcategory presented a larger difference in length (Supplementary Fig. A2d) and exon number (Supplementary Fig.A2e) with respect to their matched reference transcript than the rest of ISM, ruling out the possibility that these were fragments of initially shorter molecules. These results suggest that ISM transcripts were enriched in 5’end degradation products.

The diversity of TSS and TTS patterns in lrRNA-seq is apparent in the NEXN gene (Fig.2**f**). Although all detected multi-exon transcripts were associated with the same reference transcript model (ENST00000330010.12), they exhibited differences at their 3’ and 5’ ends that were variously supported by additional data. Two transcripts, classified as 5’ fragment ISM, displayed a loss of two exons at their 3’ end (PB.1002.4 and PB.1002.6, Fig.2f). PB.1002.4 had a shorter last exon that was supported by Quant-seq, a polyA motif, and reference annotation (i.e. shared the TTS in ENST0000440324.5) In contrast, PB.1002.6 had an extended 3’ UTR that was not supported by either the reference or Quant-seq, but did contain a known polyA motif. The NEXN gene also had three FSM isoforms that varied in their polyA sites but exhibited higher similarities at the TSS. PB.1002.1, the longest FSM, perfectly matched the reference TTS and was supported by Quant-seq. However, the remaining FSMs were 667 bp (PB.1002.2) and 1089 bp (PB.1002.3) shorter than PB.1002.1. Although polyA motifs were identified for both PB.1002.2 and PB.1002.3, these transcripts lacked Quant-seq evidence, and PB.1002.2 was flagged as a potential intrapriming artifact due to a 20-bp stretch with 90% As found immediately downstream of the TTS, which, together with the location of the polyA motif 39 bp from the 3’ end, suggested a TTS annotation of poor quality (Fig.2**f**). Notably, the degree of isoform diversity and TSS/TTS support variability exhibited by the NEXN gene was frequent in the WTC11 lrRNA-seq transcriptome estimated by Isoseq3.

In summary, the IsoSeq3 processing of the WTC11 cDNA-PacBio data presents a high level of TSS and TTS variability, which can be effectively characterized using SQANTI3 (sub)categories in combination with complementary data sources. Our analyses suggest that a combination of artifacts and true biological variability causes the observed diversity at transcript ends. These and previous SQANTI results [23] motivated the design of a comprehensive strategy for removing lrRNA-seq artifacts.

### 2.3 Characterization of the SQANTI3 Filter

SQANTI3 offers a flexible framework for removing artifacts in long-read transcriptomes, based on structural categories and quality descriptors. The Filter module can be adapted to the availability of additional data and the user’s preference for using a rules-based approach, where the selection criteria are user-defined, or a machine learning-based strategy, where a classification model is trained. In this section, we use the WTC11 dataset to demonstrate how these filtering alternatives behave in three different scenarios of increasing availability of additional data. The Low Input (LI) scenario represents a situation where only a reference annotation and matching Illumina data are available, which is typical for many studies that use long reads. In the High Input Reference (HIR) setting, additionaldata are obtained from reference databases such as refTSS and polyAsite, resembling the scenario of model species with abundant genomic resources. Finally, the High Input Sample (HIS) setting represents a case where, in addition to the Illumina reads, data such as CAGE and Quantseq have been obtained (Extended Data Table 1). To ensure comparable results between rules and ML strategies, we set the rules filter to accept transcripts models as long as their SJs, TSS, and TTS were supported by at least one source of external evidence. When running the ML filter, variables used to define the true and false transcript sets were excluded during random forest training to prevent overfitting (see Methods for details).

**Table 1.**
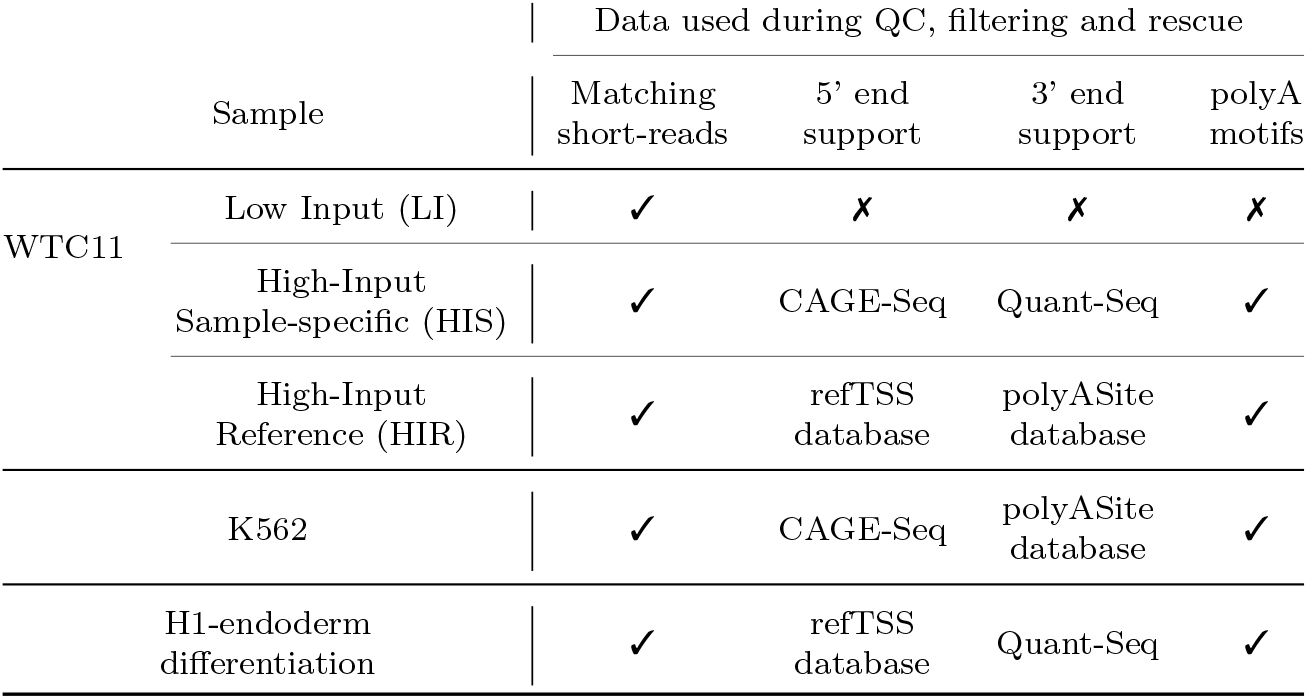
Extended Data Table 1. Data used in the different analyses.

Rules filtering with more input data increased the number of transcripts models passing the filter, as expected for the OR configuration used in its design, while the ML approach flagged a higher number of artifacts with increasing amounts of additional data (Fig.3**a**). Specifically, HIS-ML was found to yield the most stringent filtering (173,864 potential artifacts), while HIS-Rules was the most lenient (88,786 potential artifacts). Additionally, we noticed that rules-filtered transcriptomes included a significantly higher number of ISM and NNC transcripts than those obtained using the ML method (Fig.3**a**). This effect was partially mitigated in the LI scenario, where a similar structural category distribution was obtained regardless of the filtering strategy (Fig.3**a**). Interestingly, almost all transcripts flagged as artifacts using SQ3 rules were captured by the ML filter when both were run with the same orthogonal data, with the Low Input scenario being the one with the highest level of agreement (Fig.3**b**).

**Fig. 3.**
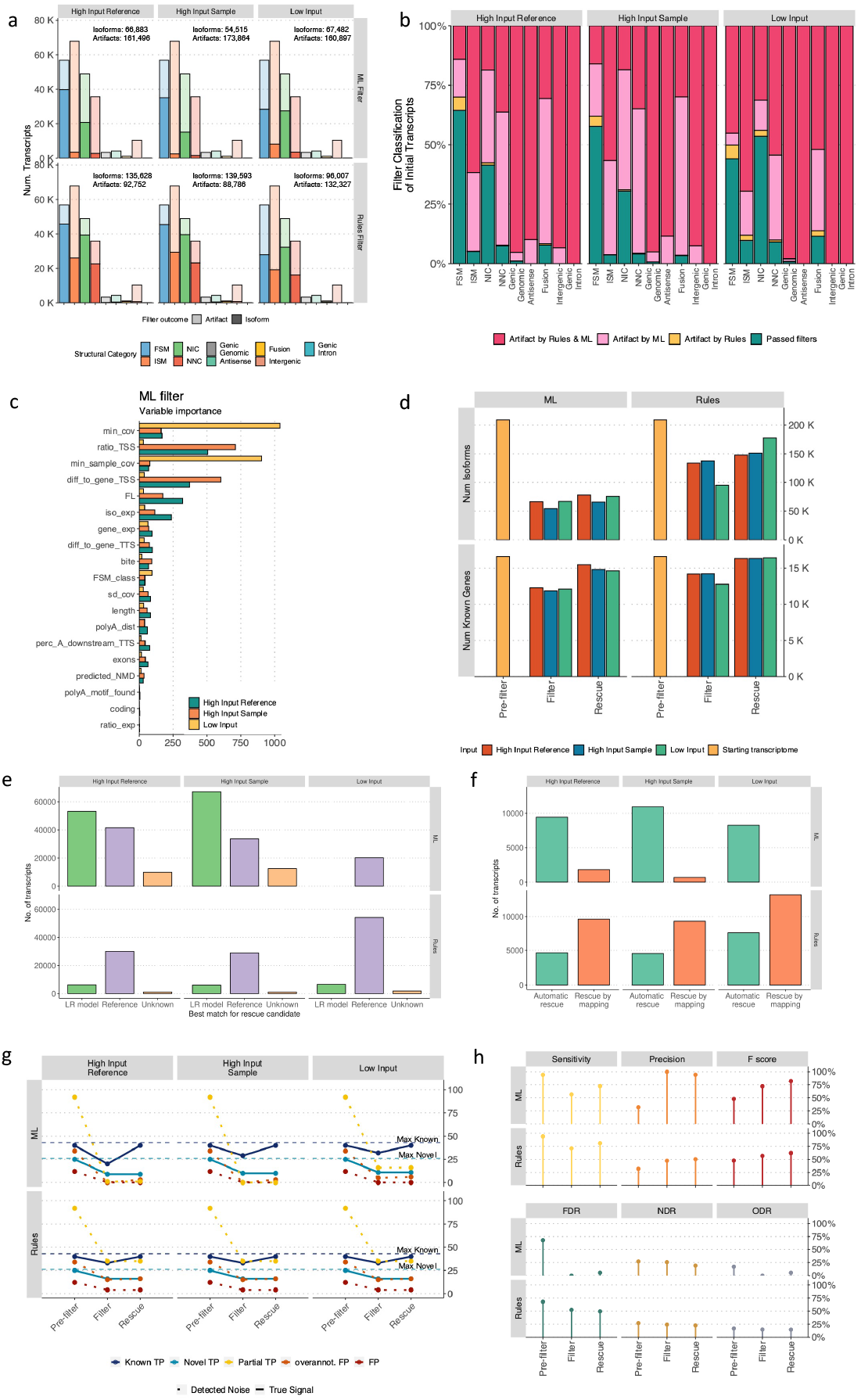
SQANTI3 curation. **a** Artifact and isoform distribution across SQANTI3 structural categories when Rules and ML filters were applied to the WTC11 transcriptome using distinct additional data. **b** Overlap in filtering outcome among filtering methods. **c** Variable importance of ML filter for different input scenarios. **d** Variation in the number of genes and isoforms after Filter and Rescue.**e** Best match between rescue candidates (transcripts filtered out) and rescue targets (reference and LR-defined transcripts that passed the filter). **f** Inclusion criteria for rescued targets. **g** Number of TP, TN, FP, and FN at SQANTI3 curation steps. **h**, Performance metrics at SQANTI3 curation steps.

Moreover, a total of 13,969 identified as artifacts in LI settings were supported within HIS and HIR configurations (Extended Data Fig.A3), suggesting that the additional TSS and TTS information is biologically relevant (e.g. PB.1195.1, Supplementary Fig.A4). This underscores the utility of these data sources for validating transcript models. Moreover, when filtering by rules, choosing sample-specific experimental data (HIS) instead of reference databases (HIR) improved the validation of novel TTSs, resulting in a decrease in the number of isoforms removed due to the absence of a polyA peak. Specifically, a total of 5,261 isoforms (69.7% of which were ISM) were retained when sample-specific data was used to inform the rules filter (Extended Data Fig.A3), supporting the notion of incompleteness in the current reference annotations.

We next examined the variables selected by the trained ML filter classifier across the different input data contexts. One advantage of this filtering strategy is that it eliminates the need for arbitrary thresholds, automating the decision to accept or reject a transcript model. However, users must still define the true positive (TP) and true negative (TN) transcript sets, which can significantly influence the filtering results. In the LI scenario, where additional evidence for TSS and TTS was lacking, the FSM Reference Match subcategory was used to define TP transcripts, while the TN set was comprised of NNC with non-canonical SJs. These groups differ in SJ quality by definition; however, this configuration did not account for sequencing incompleteness issues, such as false positive TSS/TTS. As a result, attributes related to short-read SJ coverage were the most relevant in the trained random forest model (Fig.3**c**) with minimal contributions from other variables. In contrast, TP and TN sets were defined using CAGE and Quant-seq data for the HIR and HIS scenarios to include transcripts with end variability (see Methods). Metrics such as the TSS ratio, the distance to a known TSS of the same gene, or the number of long reads supporting a transcript model were, therefore, more relevant for isoform/artifact classification **(Fig.3c)**. As a consequence, the ML-HIR and ML-HIS filters removed a greater proportion of ISM transcripts models than ML-LI, while more FSM passed the filter in these cases (Fig.3**a**). This result aligns with the SQANTI3 feature overlap analysis presented in the previous section, confirming the TSS ratio’s usefulness as a metric for validation of 5’ ends in novel transcripts and demonstrating the existence, at least in human samples, of unannotated combinations of known SJs and TTS/TSS.

In summary, we show here that the SQANTI3 Filter module is flexible enough to accommodate varying amounts of available data and can be customized to suit the user’s curation criteria. Moreover, we have found that including data supporting 3’ and 5’ ends in the validation of transcript models can significantly influence the composition of the resulting curated transcriptome. This underscores the limitations of current long-read data and emphasizes the importance of incorporating additional data when defining transcript models using long reads.

### 2.4 The SQANTI3 rescue module to enhance completeness of curated transcriptomes

SQANTI3 filtering resulted in the removal of a large number of transcript models. As a consequence, a substantial number of genes were completely excluded as none of the associated transcripts passed the filtering criteria (Fig.3**d**), despite evidence of gene expression provided by long reads mapping to these loci. To mitigate these losses, SQANTI3 now includes a Rescue module designed to identify the most likely transcript models for discarded artifacts.

When SQANTI3 Rescue was applied to ML-filtered data, an average of 10,389 isoforms and 2,884 genes were recovered from the reference transcriptome, while 36,811 transcripts and 2,638 known genes were included on average when the rescue was applied to the rules-filtered datasets (Fig.3**d**). The number of rescued genes was consistently high regardless of the input data configuration used, which highlights the consistency of the approach regarding the recovery of loci that are represented in the lrRNA-seq dataset.

To illustrate the rescue process, we discuss the ML-filtered data in a HIS scenario. The rescue algorithm identifies FSM transcript models classified as artifacts and recovers the associated reference transcript model (Methods, Supplementary Fig. A1). Artifacts from the ISM, NIC, and NNC structural categories (i.e. rescue candidates) are mapped against reference annotation and filter-passing long-read transcript models to find potential rescue targets (Methods, Supplementary Fig. A1). This resulted in the identification of multiple targets per candidate, most of which belonged to the reference transcriptome (Supplementary Fig.A5). Specifically, 59.3% of the candidates (113,320 in total) were preferentially assigned (see Methods) to a reference transcript and 29.7% matched a long read-defined isoform (Fig.3**e**). Notably, only 11% of artifacts remained unassigned to a suitable replacement transcript after the rescue process (Fig.3**e**). A large number of artifacts could therefore be matched to isoforms that were already present in the transcriptome, suggesting that an important fraction of detected artifacts may be the consequence of technical errors in library preparation and sequencing that affect correctly detected isoforms. After best match selection, targets are validated using SQANTI3 ML Filter (see Methods), which resulted in 11,599 unique candidates being selected and incorporated into the transcriptome, 94.1% of which corresponded to automatic rescues, whereas 5.9% were introduced via mapping (Fig.3**f**).

To evaluate the rescued elements, the reference transcriptome was quantified at the gene and transcript levels using short reads (see Methods). Genes for which at least one isoform passed the filter showed the highest expression values (Extended Data Fig.A6**a**), independently of the complementary data input. Genes initially removed and then rescued had consistently higher expression than those that remained excluded. Similar results were obtained when performing these analyses at the transcript level (Supplementary Fig.A7). Finally, to validate the biological importance of the rescued transcript set, we retrieved their functional scores (TRIFID scores) from the APPRIS database [34]. Known transcript models that were rescued had higher scores than those that were not rescued (Fig.A6**b**). All in all, these results demonstrate the effectiveness of the SQANTI3 rescue strategy to recover functionally-relevant known genes and transcripts that would otherwise be discarded due to limitations in the lrRNA-seq methods.

### 2.5 Validation of the SQANTI3 curation approach

To validate the ability of SQANTI3 to obtain a correctly curated transcriptome, we benefited from the usage of Spike-In RNA Variants (SIRVs) in the LRGASP WTC11 PacBio dataset. SIRVs, which are control RNAs designed to mimic splicing-related transcriptome complexity, are used to assess the accuracy of transcriptome reconstruction [35]. SIRV reads followed the same processing and curation as the rest of the WTC11 long-read data (see Methods). For this study, we modified the SIRV reference annotation to simulate two real-life scenarios that could lead to transcript model artifacts: a) an over-annotated FP scenario, in which 39 falsely annotated SIRVs-like models were included in the transcriptome, and b) a novel TP scenario, in which 26 true isoforms were removed from the reference -even though they were still present in the control RNA mix. In order to accept a transcript model as a TP, a maximum of 50bp difference at the TSS and TTS was allowed. Transcripts with a larger difference, which included ISM, were classified as Partial True Positives (PTP). Several metrics were computed based on these definitions to evaluate the performance at each stage of the curation process (see Methods).

As a result of IsoSeq3 processing, 65 out of 69 (94%) SIRVs were correctly identified, including novel TP (Fig.3**g**). However, a high number of PTP and FP were also present. After filtering, the number of FP transcripts decreased across all FP subtypes regardless of the approach and the data available, where the ML-HIS scenario proved to be the most effective with a 100% success at removing FP isoforms (Fig.3**g**). In the case of rules-HIS, 15 over-annotated FP and 35 PTP passed the filter, which can be explained by the rules filter’s reliance on the reference annotation for isoform validation. Reassuringly, however, only a reduced group of true SIRVs was cataloged as artifacts during filtering, most of which were recovered after Rescue, with the sole exception of the novel TP transcripts (Fig.3**g**). SQANTI3 Rescue was also robust to the confounding factor introduced by reference over-annotation, with only between 1 and 3 FP transcripts being selected by the algorithm.

To better understand the impact of each step of the SQANTI3 pipeline on transcriptome quality, performance metrics were computed for the HIS results (Fig.3**h**). The initial reconstruction by IsoSeq3 yielded almost perfect sensitivity (94%), while precision was much lower (32%). As previously observed, precision significantly improved after filtering, whereas sensitivity decreased slightly due to some TP transcripts being flagged as artifacts. Importantly, the F-score revealed that overall performance steadily improved after every step of the SQANTI3 curation pipeline. Specifically, a stronger F-score increase was observed when using the ML filter (82%) than when applying the rules filter (62%). The opposite trend was detected regarding the False Discovery Rate (FDR), which consistently decreased after filtering and rescue, especially when the ML filter was applied. Rules filter-based strategies, on the other hand, were less effective at removing over-annotated FP, which resulted in an almost constant Over-annotation Detection Rate (ODR), whereas ML-based approaches decreased ODR effectively. Finally, both filters yielded a constant Novel Detection Rate (NDR), showing the potential of SQANTI3-based curation to validate novel isoforms and effectively leverage the potential of long reads methods to discover new transcripts (Fig.3**h**).

In summary, the SQANTI3 pipeline was successful in reducing sequencing and transcriptome reconstruction noise while maintaining known and true novel transcript models in the final annotation. The filter strategies provided by SQANTI3 improved transcript definition accuracy, with the ML-based approach performing the best. The novel rescue strategy helped recover the majority of annotated ground truth lost during filtering, leading to a comprehensive yet highly-curated transcriptome.

### 2.6 Application of SQANTI3 pipeline to a dRNA ONT defined transcriptome

The SQANTI3 pipeline is designed to perform data-based QC and curation of transcriptomes, particularly those created using tools with high detection levels and low reference dependence, resulting in a significant proportion of novel isoforms. However, it can be applied to any *de novo* transcriptome, regardless of the sequencing technology or reconstruction pipeline used to generate it. Additionally, SQANTI3 QC can be used to identify and adjust to the algorithmic choices made during transcriptome reconstruction, detecting pipeline-specific sources of bias.

To further demonstrate its wide applicability and versatility, we used SQANTI3 to analyze a long-read transcriptome from a K562 human cell line. The transcriptome was obtained using dRNA ONT sequencing data and the ENCODE pipeline TALON [13]. This transcriptome contained 336,445 isoforms, 95% of them belonging to 51,843 known genes. Cell line-specific orthogonal data from the ENCODE site (CAGE-seq, Illumina) and reference databases (polyAsite) were used as inputs for SQANTI3 QC (see Methods). SQANTI3 identified a relatively large proportion of FSM (*∼*60%, Fig.4**a**). Of these, the vast majority (85.5%) were classified within the Reference Match subcategory (Fig.4**b**, upper panel), which involved 163,319 reference transcripts, roughly two-thirds of the entire GENCODE annotation. Further analysis revealed that one-third of all identified FSMs (33.1%) had Illumina support across all SJs and 11.4% showed expression above 1 TPM (Fig.4**b**, lower panel). These results suggest that the TALON transcriptome reconstruction process was largely guided by the reference annotation, leading to moderate support by orthogonal data.

**Fig. 4.**
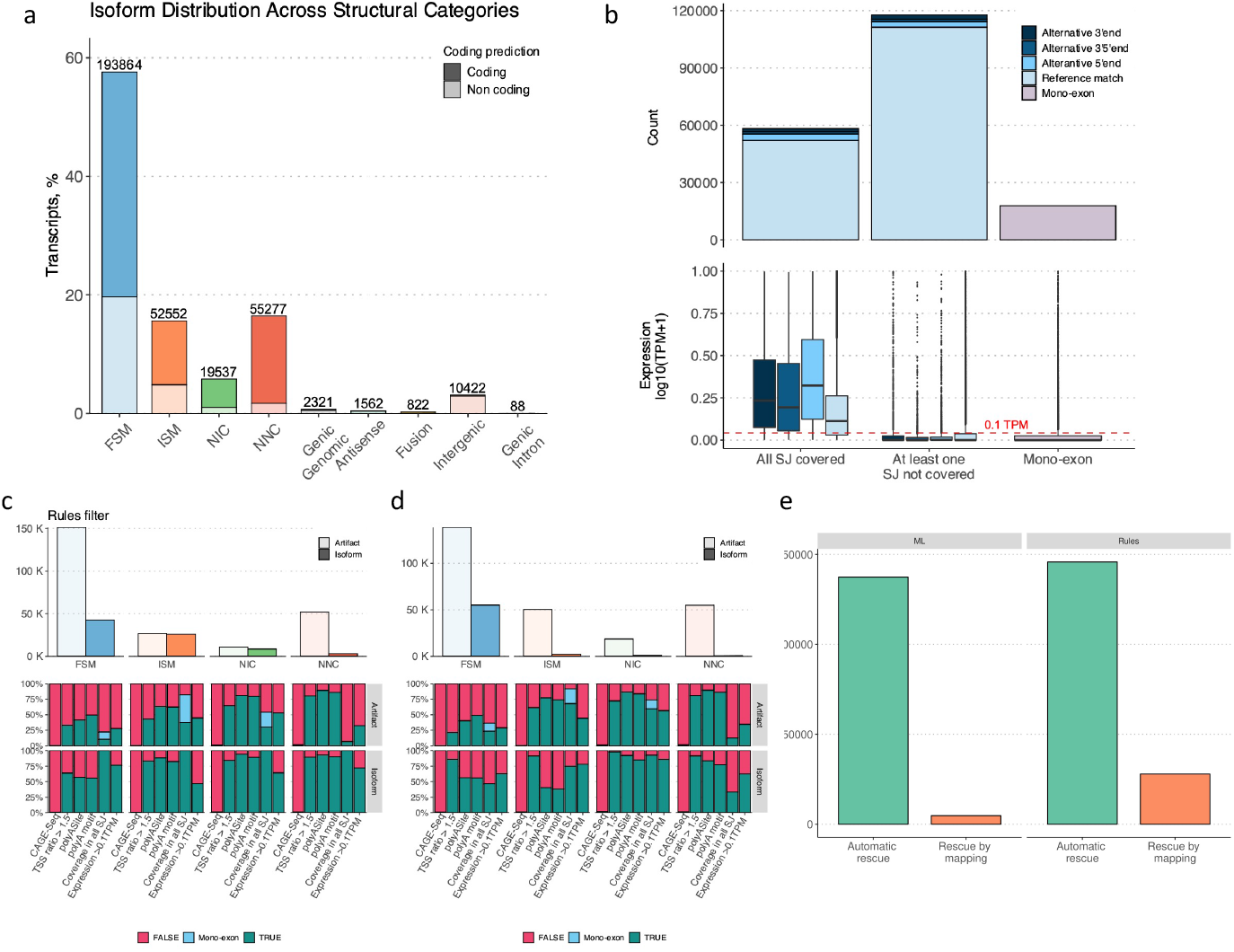
QC and filtering of the K526 dataset. **a**, Distribution of structural categories in the K562 TALON-defined transcriptome. **b**, Characterization FSM transcript models of the K562 TALON-defined transcriptome. Incidence of artifacts detected across structural categories according to SQANTI3 filter using a **c** rules or a **d** ML approach. **e** Number of transcripts incorporated into the K562 transcriptome after SQANTI3 rescue, shown by filter strategy and separated by whether they were obtained during automatic rescue or after rescue-by-mapping.

We next applied the two strategies within the SQANTI3 Filter module to the K562 dRNA ONT transcriptome. Because of the large number of FSMs with poor orthogonal support, we adjusted the rules filter to require SJ coverage of at least 3 short-reads in all structural categories, including FSM. As a result, the vast majority of FSM and NNC isoforms were flagged as potential artifacts, mainly due to their low expression values and the presence of at least one SJ lacking short-read coverage (Fig.4**c**). Moreover, the number of peaks in the available CAGE-seq dataset was low limiting the utility of this information for TSS assessment (Fig.4**c**). However, this did not prevent the validation of novel isoforms reported by TALON and after filtering by rules, the NIC and NNC categories accounted for about *∼*15% of all filter-passing transcript models, exhibiting relatively high-quality attributes (Fig.4**c**). The ML-filtered transcriptome, on the other hand, contained a remarkably high proportion of FSM and a comparatively lower number of ISM, NIC, and NNC (Fig.4**d**). We attributed this result to the reference-biased and poor SJ coverage of the TALON FSM transcripts, together with the limitations of the CAGE-seq data, both of which compromised the definition of the TP set. In fact, most filter-passing FSMs had at least one SJ lacking coverage and low short-read-based expression values (Fig.4**d**), suggesting poor support of their presence in the K562 samples. Therefore, we concluded that a tailored rules filter would be a more suitable approach in this case, as it yielded a more data-supported transcriptome. Similarly, the rescue strategy can be adapted to the characteristics of this dataset. The automatic rescue was in this case not advisable, as the high number of FSM flagged as artifacts were recovered by reference transcripts with equally poor support (Fig.4**e**). Instead, the detailed information included in the output of the rescue module can be used to define an *ad hoc* rescue criteria. In this case, we opted to only recover rescue-by-mapping isoforms, as these are supported by complementary data. The information retrieved by the SQ3 rescue module, therefore, allows for a critical evaluation of the properties of each specific transcriptome, enabling users to make decisions tailored to their dataset.

Overall, this example demonstrates the versatility of the SQANTI3 toolkit for designing a transcriptome curation strategy. Users have the ability to customize each step of the pipeline to meet their specific data needs. In the case of the TALON K562 dRNA ONT transcriptome, FSMs were in high proportion but often lacked additional evidence of expression, compromising the definition of a high-quality true positive transcript model set. Therefore, rather than using the ML approach, the rules filter was adjusted based on the information obtained through SQANTI3 QC. Additionally, the information in the output of the rescue step was used to recover only well-supported transcript models. These examples also underscore the value of complementary data in the curation process and the need for a critical assessment of each dataset.

### 2.7 SQANTI3 supports Functional Iso-Transcriptomics (FIT) analysis: an hESC differentiation example

The SQANTI3 framework offers not only quality control and curation but also the integration of IsoAnnotLite, which allows for isoform-level functional annotation. This feature facilities downstream analyses on isoform biology, for example, using the tappAS software [19]. To demonstrate this capability, we analyzed the differentiation of an H1 human embryonic stem cell (hESC) line into endoderm using replicated cDNA PacBio data generated within the LRGASP project. An experiment-specific transcriptome was created, combining the lrRNA-seq data from hESC and endoderm samples and using the IsoSeq3 pipeline (see Methods). A total of 211,107 transcripts models were initially detected, 89% of which belonged to 17,667 known genes. Data from the refTSS database and Quant-seq data, as well as same-sample short reads and a list of known human polyA motifs, were used to remove potential artifacts using the SQANTI3 ML filter (see Methods). After filtering and rescue, 65,255 transcript models from 14,541 known genes were included in the annotation (Fig.5**a**). Among them, there were 4,043 single-isoform and 10,498 multi-isoform genes (Fig.5**b**). We used stringent filtering for potential fragments and isoforms with novel SJ, resulting in a curated transcriptome that was enriched in FSMs and NICs, and had few ISM and NNC transcripts (Supplementary Fig.A8).

**Fig. 5.**
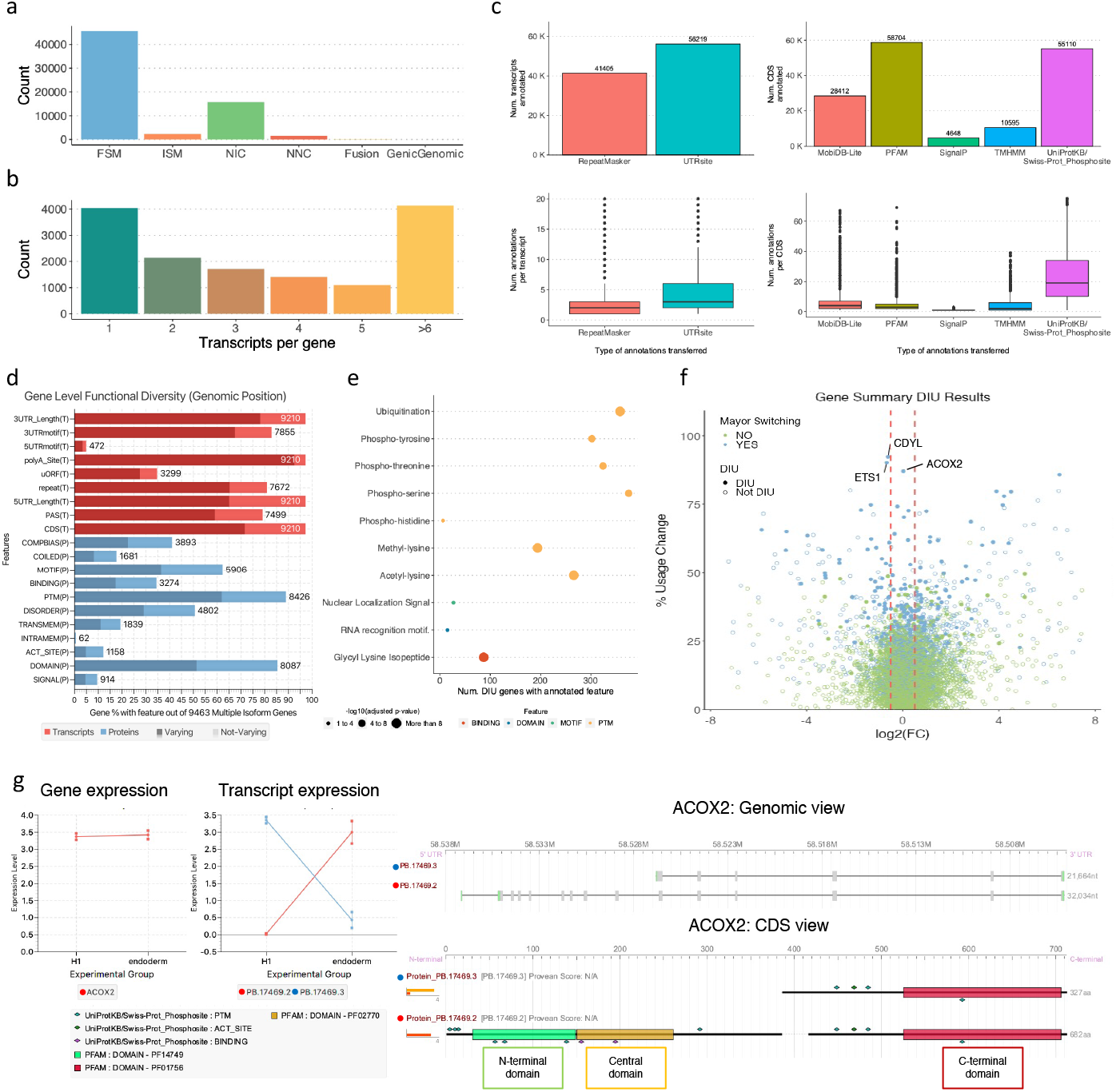
Figure 5: Functional IsoTranscriptomics Analysis of the H1 hESC to endoderm differentiation. **a**, Isoform distribution across structural categories of the long-read defined H1 transcriptome. **b and c**, Functional annotation success for transcripts and CDS. **d**, Annotation of multi-transcript genes **e**, Functional Enrichment Analysis of annotated features in those genes with Differential Isoform Usage compared to Deferentially Expressed genes. **f**, Percentage of Isoform Usage Change of identified genes (y-axis) compared to their expression fold-change in expression (x-axis). **g**, Expression at the gene and transcript levels for ACOX2. **h**, Genomic and CDS views of the identified isoforms for the ACOX2 gene, including functional features annotated with IsoAnnotLite.

Next, functional annotations predicted for the reference transcriptome [19] including 9 transcript-level and 11 protein-level feature types were transferred to the long-read defined transcript models using IsoAnnotLite (see Methods). The vast majority of isoforms (82.7%) were functionally annotated with at least one functional element, being UTRsite motifs [36] (86.2% of transcripts annotated) and PFAM domains (90% of predicted CDS annotated) the most frequently transferred terms (Fig.5**c**). Short read-based quantification of isoform expression was then performed for each sample and replicate, and both isoform quantification and annotation data were used as input for tappAS analysis of H1-hESC versus endoderm differences. Using tappAS’ Functional Diversity Analysis (FDA, [19]), we observed that the majority of multi-isoform genes showed changes in the inclusion of transcript (50%) and protein (70%) features in at least one isoform, indicating that a high amount of predicted functional features were associated with potential changes in isoform functionality (Fig.5**d**).

A total of 4,247 (29.25%) genes and 21,489 (36,2%) transcripts were found to be significantly Deferentially Expressed (DE, FDR *<* 0.05) when comparing hESC and endoderm. This analysis recapitulated the behavior of known markers of differentiation [37], namely the downregulation of OCT4, SOX2, and NANOG in endoderm and the upregulation of FOXA2, SOX17, and GATA4 (Supplementary Fig. A9). To understand the contribution of Alternative Splicing (AS) to this process, the tappAŚs Differential Isoform Usage (DIU) analysis was applied. We detected 450 genes with significant DIU (FDR *<* 0.05), revealing a weaker contribution of AS to the divergences between both differentiation stages than that of DE genes. These genes were enriched in ubiquitination sites, RNA recognition domains, and Nuclear Localization Signals (adjusted p-value *<* 0.05, Fig.5**e**), suggesting a splicing-based regulation of the inclusion of these functional elements.

To identify genes in which alternative splicing was likely to be the main regulatory mechanism, we selected candidates that exhibited significant DIU (FDR *<* 0.05) but no differentiation-induced changes in gene-level expression between the two cell stages. The ACOX2 gene was found among the DIU genes with the highest isoform usage change (Fig.5**f**) while also showing no differential expression (Fig.5**g**). This gene encodes the Acyl-CoA Oxidase 2 enzyme, responsible for fatty acid degradation in the peroxisome [38]. At the structural level, the major isoform in the H1 cell line lacked the first 9 exons, which resulted in the H1-specific loss of the N-terminal and central peroxisomal Acyl-CoA oxidase protein domains (Fig.5**h**). Both domains were deferentially excluded in the H1-hESC sample (DFI adjusted p-value *<* 0.05). Upon differentiation into endoderm, the major isoform switch caused the long isoform to be almost uniquely expressed, leading to a condition-specific inclusion of the three PFAM domains necessary for full functionality.

The ETS1 transcription factor was another interesting example of development-related changes in the transcriptomic landscape that would be masked if the gene-level expression were considered exclusively. ETS1 is a gene known to be involved in cell development [39] and did not exhibit significant variation in the H1 transition H1 from hESC to endoderm. However, significant changes in transcription start site (TSS) usage were detected when analyzing isoform expression differences between the two cell types (DIU p-value *<* 0.05), which ultimately changed coding sequence (CDS) length (Extended Data Fig.A10**a-c**). The shorter protein (UniProt ID:P14921-1), highly expressed in endoderm, includes two Lys-Gly isopeptides that interact with SUMO2, while the longer isoform (UniProt ID:P14921-3), highly expressed in H1, showed exclusion of these sites (Extended Data Fig.A10**d**). Our analysis suggests that this isoform switch in the ETS1 gene may involve the gain of the ability to bind a post-transcriptional modifier during differentiation, which can constitute a splicing-mediated regulation of the role of this transcription factor.

By using long-read RNA sequencing (lrRNA-Seq) to generate the transcriptome, we were also able to identify functionally-relevant genes that undergo DIU involving isoforms not described in the reference annotation. For instance, the Chromodomain Y-like (CDYL) gene, which encodes a chromatin reader protein involved in gene activation and repression via chromatin change [40], was found to have up to five distinct isoforms identified by lrRNA-seq. Two of these isoforms were NIC by the combination of known splice sites and differed from each other only at the 3’UTR (Extended Data Fig.A11**b**). The remaining 3 isoforms were FSMs of ENST00000397588.8 (Reference Match and Alternative 3’ end) and ENST00000472453.5 (Alternative 3’ end). The most highly expressed isoforms in H1 and endoderm were PB.21000.1 (longest NIC) and PB.21000.6 (FSM Reference Match), respectively (Extended Data Fig.A11**a-c**). The usage of an alternative first exon by the novel isoform PB.21000.1 in H1 resulted in the exclusion of the chromodomain at the N-terminus (DFI adjusted p-value *<* 0.05), which was re-gained in endoderm after the isoform switch with the domain-including FSM PB.21000.6 (Extended Data Fig.A11**d**). This isoform switch may have implications for the activity of CDYL during cell differentiation, possibly involving a splicing-regulated increase in the ability of this protein to change chromatin architecture. These results emphasize the significance of augmenting the reference annotation with transcript models acquired from lrRNA-seq. Moreover, they demonstrate that when combined with functional annotation, lrRNA-seq has the potential to detect novel isoform switches of potential biological relevance.

## 3 Discussion

The development of long-read sequencing technologies has created new avenues for analyzing the transcriptome but also presents new challenges as both the sequencing platforms and the analysis methods may introduce errors when inferring transcript models from the data.

SQANTI3 addresses these QC issues by defining useful structural categories to describe the data, computing quality parameters, and incorporating various complementary data sources that are informative of transcript structural elements. These features enable the comprehensive curation of lrRNA-seq transcriptomes. Some of the additional data types that SQANTI3 is able to process, such as CAGE-seq and Quant-seq, are costly and difficult to obtain on a general basis. Aware of this limitation, SQANTI3 introduced the TSS ratio metric based on short reads and identifies polyA motifs close to the end of the transcript for the same purpose, providing a cheaper alternative for curation. Additionally, one potential risk of applying the QC criteria is the removal of poorly supported transcript models that still represent a true transcriptional signal. SQANTI3 offers an alternative to rescue this signal by providing reference transcripts as a confident replacement, which has been shown to increase the accuracy of the resulting transcriptomes. Moreover, SQANTI3 is able to highlight potential biases associated with the transcript reconstruction tool (i.e. IsoSeq3, TALON) and establish a tailored strategy to reduce their impact. The SQANTI3 analysis shown here confirms that novel combinations of 3’ and 5’ ends with intron chains are still to be discovered, even in well-annotated organisms such as human and mouse, and that many novel transcripts can be found with considerable support. This implies that the generation of sample-specific transcriptomes enabled by long-read sequencing is recommended for studies where alternative isoform expression is a goal. The H1-hESC example presented in the paper highlights the significance of lrRNA-seq, revealing novel isoforms might be part of differentiation programs and facilitating the characterization of the impact of AS on cellular differentiation. SQANTI3 adds to this discovery of novel biology the unique capability of providing functional annotation with isoform resolution, whereby predictions on the functional consequences of the alternative isoform usage can be made.

In conclusion, SQANTI3 is a comprehensive tool for the curation, structural, and functional annotation of lrRNA-seq-derived transcriptome datasets. It can integrate orthogonal data to improve the accuracy of transcript models and provides a range of filtering options to accommodate different research goals. With the ability to discover novel transcripts and provide functional annotation, SQANTI3 is an important tool for researchers studying the transcriptome.

## 4 Methods

### 4.1 Novel features in SQANTI3

#### 4.1.1 SQANTI3 Quality Control module

SQANTI3 is the latest version of the SQANTI tool https://github.com/ ConesaLab/SQANTI, originally released in 2018 [23]. All results generated in this manuscript were obtained using version 5.0 of SQANTI3. The software is publicly available on GitHub https://github.com/ConesaLab/SQANTI3, together with a comprehensive wiki in which terms, data processing steps, input and output files are described.

The Quality Control module is the cornerstone of the SQANTI3 pipeline. It is designed to characterize transcriptomes built using lrRNA-seq data and make QC decisions according to the purpose of the study. Combining Python and R scripts, SQANTI3 QC compares the *de novo* transcriptome against a reference annotation and outputs an easy-to-explore*classification* file. This file is the source of the graphical output, is used as input for the Filter module, and is generated for the reference annotation to run SQANTI3 Rescue. For clarity, improvements and modifications with respect to SQANTI are outlined in this section.

##### New subcategories for Full Splice (FSM) and Incomplete Splice Matches (ISM)

SQANTI3 QC expands the characterization of the variability at the ends of the transcript models by creating subcategories for reference-associated transcript categories, i.e. FSM and ISM.

FSM can be directly associated with known transcript models based on their SJs, but differences at their ends are not negligible. When differences are small, meaning that the FSM transcript ends match the reference with a difference *≤*50bp upstream or downstream from the annotated TSS and TTS, the transcript models are assigned to the Reference Match subcategory. Conversely, transcripts with *≥*50bp distance to the annotated TSS or TTS are considered ”alternative”, and annotated to one of the three additional subcategories: Alternative 5’-end, Alternative 3’-end, and Alternative 5’/3’-ends. For ISM, which are fragments of annotated SJ combinations, subcategories are defined based on the missing fraction of the known transcript. ISM can therefore be classified as 3’ Fragments, 5’ Fragments, and Internal Fragments depending on whether they lost exons at in their 5’, 3’, or both ends. An additional subcategory, ISM with Intron Retention (IR), was defined to classify transcripts for which the loss of an SJ in the reference is due to an IR event. Finally, the mono-exon subcategory was included for both structural categories since, despite their lack of SJs, they can be associated with known transcripts by overlap. Specifically, if a mono-exon transcript overlaps a mono-exonic reference, it will be cataloged as FSM, whereas if the reference is multi-exonic and the query sequence lies within the boundaries of an annotated exon, it will be considered an ISM. NIC and NNC categories remain the same as they were defined in SQANTI[23].

##### Integration of evidence around 5’/3’-ends

In addition to short and long-read data for coverage and expression based-validation, SQANTI3 QC can make use of additional data to generate metrics related to TSS and TTS support, namely CAGE, Quant-Seq data or other region-based sources of information providing TSS/TTS evidence. Specifically, the QC module accepts BED files including the genomic coordinates of the specific regions, e.g. CAGE peaks called, as input. The overlap between each TSS and TTS reported in the lrRNA-Seq transcriptome and the supplied regions is verified, flagging cases where transcript ends fall within a peak. Additionally, the distances between the TSS/TTS and the middle point of the closest peak are computed. Only peaks upstream of the TSS or downstream of the TTS are interrogated for this purpose.

##### Processing of short read data

To facilitate the integration of matching short-read data, SQANTI3 has been upgraded to run STAR [41] and Kallisto [42] internally for mapping and quantification purposes. Whereas SQANTI needed pre-computed mapping and quantification files, the QC module now accepts raw Illumina data as FASTQ files and can automatically generate short read-related quality metrics.

First, a genome index is created and short reads mapped short reads to identify individual SJs using STAR. The reference annotation is not used in this process to make SJ identification completely independent from prior annotations. Mapping parameters used in STAR are adapted from the ENCODE-DCC protocol for RNA-seq (https://github.com/ENCODE-DCC/ rna-seq-pipeline/), however, in order to improve the detection and quantification of novel SJs, the --twopassMode option is activated [30]. After running STAR, a *SJ.out.tab* and a BAM file are generated for each replicate. The *SJ.out.tab* file contains the quantification of long-read-defined SJs by shortread, whereas the BAM file is used to calculate the novel TSS ratio metric.

Taking the genomic position of all the long-read-defined TSS in the query annotation, two BED files including the 100bp regions downstream (inside the first exon) and upstream (outside the first exon) of the TSS are created. Using BEDTools [43], short read coverage across both genomic segments is measured and the TSS ratio is calculated with the following formula:

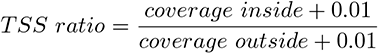

For each TSS, SQANTI3 QC computes as many ratio values as provided short-read sequencing replicates, though only the highest ratio is finally retained.

For transcript model quantification, SQANTI3 runs Kallisto internally when short-read paired-end data is provided. First, an index is built using the transcript sequences extracted from the query annotation and the reference genome. Then, short reads are pseudoaligned to these genome-corrected sequences to quantify them using --bootstrap-samples 100 as the only non-default parameter.

Further details regarding the parameters used to run STAR and Kallisto within SQANTI3 QC can be found in the Supplementary Methods section.

#### 4.1.2 SQANTI3 Filter module

##### Machine learning-based filter

SQANTI3 substantially improves the SQANTI machine learning-based filter (ML filter) [23]. Briefly, the filter is based on the training of a random forest classifier to calculate the probability that a given transcript is an isoform or an artifact. The training process is based on SQANTI3 QC attributes and on the selection of true positive (TP) and true negative (TN) isoform sets that will show differences in these attributes.

IN SQANTI3, the definition of TP and TN sets has been modified to allow either user-defined transcript lists for model training or using an automated selection of TP and TN transcripts based on SQANTI3 categories and subcategories when no TP and TN lists are supplied. By default, NNC non-canonical isoforms are taken as TN, whereas the reference match (RM) subcategory is used as TP, as these transcript models have both reference-supported junctions and ends. In addition, the size of these sets is now automatically balanced, downsizing the largest to match the number of transcripts in the smallest training list. The maximum training set size can be adjusted by the user to alleviate the computational burden of model training, even in cases where TP and TN are automatically defined. Relative to filtering criteria, the SQANTI3 implementation of the ML filter has been designed to be more stringent on the isoform than on the artifact condition, requiring that the probability that the transcript is a true isoform be *≥*0.7 by default. This threshold, nevertheless, can be modified by the user. Finally, the ML filter now allows users to force the inclusion or exclusion of some specific isoform groups. For FSM transcripts, users may indicate that all FSM should be included as true isoforms in the filtered transcriptome. In addition, since mono-exonic transcripts are not evaluated during the ML filter due to the lack of junction-related attributes, these can be automatically removed from the transcriptome if desired; otherwise, they will not be subjected to filtering.

A key improvement to the ML filter in SQANTI3 is the usage of the novel TSS and TTS validation metrics and additional data incorporated during QC for classifier training, namely CAGE-seq peaks, the short read-based TSS ratio metric, polyA motif information and Quant-seq peaks. As a result, the filter can now detect artifacts belonging to the FSM and ISM categories, which were automatically flagged as true isoforms in the original release. By default, all available variables in the SQANTI3 classification file are used for model training, except those related to genome structure (e.g. chromosome/strand), to associated reference transcript or gene, and SQANTI categories and subcategories. However, the new ML filter includes the possibility to further exclude variables, a feature designed to prevent overfitting in cases where one or more of these variables have served as criteria for TP and TN set definition. Additional details, including how to run the ML filter, are available at the SQANTI3 wiki site https://github.com/ConesaLab/SQANTI3/wiki/ Running-SQANTI3-filter#ml.

##### Rules filter

To make the Filter module more flexible and accommodate cases when the definition of TP and TN sets is not possible, a rules-based strategy for artifact removal has also been included in SQANTI3. The strategy is based on the usage of a JSON file [44] in which the characteristics that make an isoform reliable are specified.

The JSON file is structured in two different levels of hierarchy: rules and requisites. A rule is made of one or more requisites, all of which must be fulfilled for an entry to be considered a true transcript. This means that requisites will be evaluated as AND in terms of logical operators. If different rules (i.e. sets of requisites) are defined for the same structural category, they will be treated independently from one another. In that case, to pass the filter, transcripts will need to pass at least one of these independent rules, meaning that rules will be evaluated as OR in terms of logical operators. Rules can be set for any numeric or categorical column in the classification file. Numeric value thresholds can be defined as a (closed) interval of accepted values or by just setting the lower limit. For categorical QC attributes, users may define one or more acceptable levels. More details and examples of how to correctly define the rules are available at the SQANTI3 wiki site https://github.com/ConesaLab/SQANTI3/wiki/Running-SQANTI3-filter#rules.

The default filter includes two sets of rules, one for FSM transcripts and one for the remaining categories. FSMs will be removed solely based on intra-primming, i.e. if they have *≥*60% of As in the sequence immediately downstream to the TTS of the transcripts. Transcripts from other categories will require to be negative for intra-primming and RT-switching, as well as to have all of their junctions supported by at least 3 short reads or to have only canonical junctions. However, JSON-based rule definition allows users to apply *ad hoc* filtering criteria to each SQANTI3 category, which may involve thresholds for multiple QC attributes of interest and take advantage of any orthogonal data available to obtain a transcriptome adapted to each research situation.

#### 4.1.3 SQANTI3 Rescue module

The SQANTI3 rescue algorithm was conceived to be run after removing potential artifacts from the transcriptome using the SQ3 filter module to identify a *bona fide* transcript model for the discarded transcripts. The goal is to avoid the loss of transcripts and genes that constitute part of the transcriptional signal, but for which a correct transcript model could not be generated when processing the long-read data. In practice, this is done by selecting a replacement transcript from the reference that is ultimately added to the set of long read-defined, filter-passing isoforms to generate an expanded, final version of the transcriptome. This strategy is based on two principles: consistent quality, meaning that rescued transcript models should meet the QC criteria of the filtering settings used to call isoforms, and non-redundancy, meaning that when the identified replacement transcript for a given artifact is already part of the filtered transcriptome, no transcript model is added. The SQ3 rescue algorithm operates in four steps.

The first step in this module, hereby referred to as automatic rescue, applies to FSM artifacts, which may originate when TSS/TTS validation could not be performed with sufficient confidence. In this case, reference transcripts associated with at least one removed FSM are automatically added to the transcriptome. When multiple removed FSM have the same associated reference transcript but different TSS, the reference sequencing is added only once. Artifact transcripts from the ISM, NIC, and NNC categories are considered as *rescue candidates* and will continue to be analyzed by the rescue pipeline. ISM artifacts are only considered if they do not have an FSM counterpart associated with the same reference transcript.

A group of *rescue targets*, i.e. potential replacement transcripts for the *rescue candidates*, is next defined. This includes all long read-defined and reference isoforms for which at least one same-gene rescue target was found. Matches between each rescue target and its same-gene candidates are next found by mapping candidate sequences to targets, a process known as rescue-by-mapping. To achieve this, minimap2 [45] is run in the long-read alignment mode using the -a parameter, combined with the -x map-hifi (i.e. highfidelity read alignment) preset option to reflect the accuracy of the processed transcript sequences. Secondary alignments are allowed and set to the default number of 6 to allow multiple mapping hits to be reported per candidate. This process yields a series of alignments that pair each rescue candidate to multiple possible targets, pairs that are hereby referred to as mapping hits.

To ensure that reference transcriptome targets included in the rescue process comply with the same quality requirements as long-read targets and minimize the risk of retrieving non-sample-specific transcripts, SQANTI3 QC and Filter modules are run on the reference transcriptome supplying the same additional data and quality criteria as used for the long-reads transcript models. The rescue-by-mapping process is completed by applying a series of criteria to select suitable targets for inclusion in the final transcriptome. First, mapping hits are disregarded if the rescue target did not pass the filter, be it ML or rules. When multiple mapping hits per rescue candidate exist that passed the filter in the ML strategy, the target transcript with the highest ML filter probability (long-read or reference) will be selected for rescue. If multiple hits pass the rules filter, all targets will be considered, since there are no further refinement criteria available in this case. If the best match target is a long read-defined isoform, which is already included in the transcriptome, no further action is performed, as the artifact is already represented in the dataset. The remaining reference targets are then evaluated for potential redundancy against the curated transcriptome and only added if not already present.

#### 4.1.4 IsoAnnotLite and tappAS integration

IsoAnnotLite is a Python script for the transference of isoform-level functional feature annotations from an existing tappAS annotation GFF3 file to long-read-defined transcripts processed using SQANTI3 QC. During this process, long-read-defined transcripts, i.e. *feature acceptors*, receive annotations from transcripts that are already annotated with functional information and therefore act as *feature donors*. The resulting GFF3 file is compatible with the tappAS software for isoform-level functional analysis. In case a reference GFF3 annotation file is not provided, the output transcriptome will be formatted into a tappAS-compatible GFF3 file including only structural information, enabling quantitative but not functional analysis within tappAS.

The script uses the classification, junction, and GTF file generated by SQANTI3 QC, and can be run simultaneously as the QC script by supplying the --isoAnnotLite flag and providing a tappAS GFF3 functional annotation file via the --gff3 argument. IsoAnnotLite is executed with -novel as the only non-default parameter to force all transcripts to be treated as novel transcripts, meaning that each long-read isoform or feature acceptor is annotated using functional information from all feature donors belonging to the same gene in the reference GFF3. When the -novel argument is not supplied, the non-novel acceptor transcript will only receive annotations from donor transcripts with matching IDs. More information on IsoAnnotLite parameters and their effect on the annotation process is available in the SQANTI3 documentation: https://github.com/ConesaLab/SQANTI3/wiki/IsoAnnotLite.

The IsoAnnotLite algorithm includes the following steps. First, positional information for feature acceptors, i.e. transcripts in the lrRNA-seq transcriptome, is converted to genome positions using the information in the SQANTI3 QC output. Similarly, transcript-level functional feature positions from the donor transcripts are transformed into genomic coordinates using the information in the reference GFF3 file. Next, functional features are transferred across transcript models by matching genomic positions, i.e. features from the donor transcripts whose genomic positions span a feature acceptor will be annotated as belonging to that transcript. It should be noted that different transfer rules have been implemented depending on the type of feature that is being handled. To transfer UTR features, genomic feature positions must be inside the transcript’s exons and outside its CDS region. For CDS transcript features (namely transcript-level features situated within the coding region), 1) the feature must be contained within the acceptor transcript’s exons as well as inside the CDS region; and 2) if a feature has start and end positions situated in different exons, the end and the start of the exons for the donor and acceptor transcripts must be the same in order for IsoAnnotLite to transfer the feature. For protein features donor and acceptor transcripts are first verified to be coding and have the same CDS. If all CDS exons are the same for both transcripts, all protein features are automatically transferred. If not, IsoAnnotLite requires the genomic positions of at least one CDS exon to be a partial match, that is, for the feature donor and acceptor to share part of one exon in the transcript’s CDS. If at least one CDS genomic region overlaps between both transcripts, IsoAnnotLite checks for protein features that fall within that region and can therefore be transferred. For gene-level characteristics (e.g. Gene Ontology terms), information is always transferred across matching gene IDs. Finally, IsoAnnotLite verifies whether the same feature has been transferred from multiple donor transcripts to the same acceptor, and performs the removal of duplicated annotations.

### 4.2 Data

#### 4.2.1 WTC11 dataset

The WTC11 cell line is an induced pluripotent stem cell (iPSC) line derived from human fibroblasts, often used as a model for cell differentiation [46]. The data used in this paper was generated within the LRGASP project [31], where this cell line was deeply sequenced using different technologies. We only used cDNA-PacBio data for reconstructing transcripts models. Raw data used in this manuscript (subreads) is accessible through the ENCODE database, under experiment accession ENCSR507JOF. Additionally, raw short-read data from the same samples were retrieved from ENCODE experiment accession ENCSR673UKZ. In both cases, sequenced RNA samples included Lexogen’s Spike-In RNA Variants SIRV-Set 4 (Cat. No. 141). This included 69 short SIRVs, 15 long SIRVs, and 92 mono-exonic External RNA Controls Consortium (ERCC). Short SIRVs (191-2.5kb) are designed to reproduce different splicing patterns with respect to their reference gene, creating a multi-isoform scenario. In contrast, long SIRVs (4-12kb) do not present splicing. The 69 short SIRVs resemble -multi-exonic genes with alternatively spliced isoforms- and were used as ground truth to evaluate performance after each step in the SQANTI3 pipeline.

CAGE-Seq data for the WTC11 cell line was obtained from LRGASP (GEO accession GSE185917), while Quant-Seq data was downloaded from ENCODE experiment ENCSR322MWL. Reads were processed to obtain a collection of sample-specific peaks, which we filtered to only include peaks found in at least two replicates, resulting in 46,722 CAGE-Seq and 45,813 Quant-Seq peaks. Reference annotations of TSS and TTS were obtained from the Reference Transcription Starting Sites [32] (refTSS, v3.1) database and from the polyA- Site database [47] (v2.0), respectively. A list of common human polyA motifs was obtained from [33].

#### 4.2.2 K562 dataset

The K562 commercial cell line is derived from lymphoblasts from a chronic myelogenous leukemia patient and it is used as a model for immunological research [48]. For the purpose of this study, a lrRNA-Seq K562 dataset obtained using direct RNA (dRNA) Oxford Nanopore Sequencing was retrieved from ENCODE accession ENCSR917JIA. Specifically, we downloaded the reconstructed long-read transcriptome obtained using the TALON pipeline [13], which was available under ENCODE accession ENCFF584GRG.

Additional data were retrieved from a variety of experiments involving the same cell line, available at the ENCODE database. Short-read, single-end Illumina data from two K562 samples were obtained from ENCODE accession ENCSR792OIJ. CAGE-Seq data was downloaded from ENCODE accession ENCSR000CJN and included 9,248 peaks. Given the unavailability of K562 Quant-Seq datasets, the reference database described in the previous section (polyASite v2.0, human) was used for TTS validation, together with the same list of human polyA motifs. To obtain the read counts associated with each transcript model, we parsed the information from the transcript quantification file under the ENCODE accession ENCFF668BLB. All these data were used as input for SQANTI3 QC.

#### 4.2.3 H1-endoderm differentiation dataset

To demonstrate the functional significance of SQANTI3 curation, we selected a human Embryonic Stem Cell (H1-hESC) dataset in which H1 cells had been differentiated *in vitro* into human Definitive Endodermal (DE) cells.

These two cell cultures were sequenced using both Illumina short-read sequencing and PacBio lrRNA-Seq as part of the LRGASP project. Long-read data for these samples is available at ENCODE accessions ENCSR271KEJ (H1) and ENCSR127HKN (DE). Long reads from all samples and replicates were pooled to reconstruct a single transcriptome using IsoSeq3 (see Supplementary Methods). Illumina data was retrieved from ENCODE accessions ENCSR588EJX (H1) and ENCSR266XAJ (DE). The refTSS database was to retrieve human TSS annotations, while TTS support was computed by combining a human polyA motif list (see above) and 52,558 peaks from sample-specific Quant-Seq data supplied by the LRGASP consortium (ENCODE accession: ENCSR198UNH).

### 4.3 Data processing

#### 4.3.1 Transcriptome reconstruction with IsoSeq3

The PacBio cDNA lrRNA-Seq datasets used in the present study, i.e. WTC11 and H1-DE, were processed using the IsoSeq3 software (v3.4.0) for *de novo* long-read transcriptome reconstruction, provided by PacBio (https://isoseq.how/). We followed the recommended pipeline by PacBio to build a transcriptome starting from subreads:

1. For each replicate in any given dataset, the ccs function was run using default parameters and setting --min-rq 0.9 to define the minimum predicted accuracy.
2. The lima function was next run using the --peak-guess and --isoseq arguments and default parameters. This allows the identification of primers and the removal of chimeric reads.
3. Primer sequences were then trimmed using IsoSeq3 refine, with the --require-polya argument to only keep reads where both primers and a polyA tail were identified i.e. Full-Length Non-Chimeric (FLNC) reads.
4. Replicates (and samples) were pooled together for the IsoSeq3 cluster step.
5. Transcript collapse was then performed to minimize transcript model redundancy using cDNA Cupcake (https://github.com/Magdoll/cDNA Cupcake/). After mapping transcripts to the genome with minimap2 [45], the collapse isoforms by sam.py script was run using the --dun-merge-5-shorter flag to prevent the removal of alternative TSS.

For further details on the exact code used to run IsoSeq3, see Supplementary Methods.

#### 4.3.2 Running SQANTI3 QC

SQANTI3 QC was run using the human GENCODE annotation (v39) and default parameters for all three datasets. The different types and sources of orthogonal data used in each case are reported in Extended Data Table 1.

In the case of the K562 cell line, available short-read RNA-seq data was single-end, which is not supported by SQANTI3 QC short-read processing. In this case, Kallisto was run independently and predicted transcript abundances (TPMs) used to build an expression matrix, which was supplied to SQANTI3 QC via --expression argument.

#### 4.3.3 Running SQANTI3 filters

##### Rules filter

For the WTC11 dataset, the Rules Filter was defined as follows. The 5’-ends were considered valid if: 1) they overlapped a CAGE-Seq peak or an annotated refTSS site; 2) the distance to any other annotated TSS in the same gene was less than 50bp; or 3) they had a TSS ratio 1.5. Similarly, 3’-ends were accepted if: 1) they were supported by Quant-Seq data or by polyASite annotation, 2) the distance to any other annotated TTS was less than 50bp, or 3) there was a canonical polyA motif close to the TTS. FSM and ISM were required to have support in both their 5’ and 3’-ends to pass the filter. For the rest of the transcript models, it was required that all SJ were supported by at least three short-reads or were canonical junctions. Additionally, isoforms were filtered out if labeled as an intrapriming artifact (60% A’s in the 20bp downstream of the reported TTS at the genomic level), if one or more SJs were flagged as generated by RT-Switching or if they were mono-exonic.

For the K562 dataset rules were modified to require 5’ and 3’-end validation (as defined above) as well as SJ coverage by at least 3 short reads in all junctions. Similarly to the WTC11 rules, isoforms were removed if mono-exonic or flagged as intrapriming or RT-Switching.

##### ML filter

The definition of True Positive (TP) and True Negative (TN) transcript model sets is critical for the performance of the ML filter. When TSS/TTS orthogonal data was available (WTC11 HIS and HIR scenarios, H1 differentiation experiment and K562 dataset), the TP set was defined using FSM multi-exonic isoforms with CAGE support at the 5’ end, Quant-Seq support at the 3’ end and canonical SJ. Meanwhile, the TN set consisted of non-FSM multi-exonic transcripts lacking support in at least one of their ends or presented a noncanonical SJ. In total, 3000 transcript models showing these properties were randomly sampled for each set. Importantly, the attributes used to define TP and TN sets, i.e. CAGE and Quant-Seq support and canonical status of SJs, were removed from the set of variables used for model training in order to prevent overfitting. To achieve this, classification column names were supplied to the ”–remove columns” option in the SQANTI3 ML filter. Finally, given the unavailability of TSS/TTS orthogonal data in the WTC11 LI scenario, built-in TP and TN sets were used, namely FSM in the Reference Match subcategory were considered TP, while NNC isoforms with at least one non-canonical SJ were defined as TN.

#### 4.3.4 Running SQANTI3 rescue

In order to perform the rescue step, the reference annotation was first characterized using SQANTI3 QC and the same additional data employed during lrRNA-seq transcriptome QC. However, since SQANTI3 ignores by default reference transcripts shorter than 200bp to avoid spurious matches, to appropriately characterize the reference, we set --min ref len 0 in SQANTI3 QC to avoid the otherwise misclassification of short transcripts.

Next, SQANTI3 rescue was run using the SQANTI3 QC classification file from the reference and the SQANTI3 filter output obtained after filtering the long read-defined transcriptome as input. In addition, for post-ML filter rescue runs, we supplied the pre-trained Random Forest classifier used for transcriptome filtering. For post-rules filter rescue runs, the JSON file containing the rules and requisites used upon filtering was supplied as input to SQANTI3 rescue.

#### 4.3.5 Running IsoAnnotLite and tappAS

For the functional annotation of the H1-DE dataset, the SQANTI3 QC script was re-run using the post-rescue transcriptome as input and the --isoAnnotLite and --gff3 arguments. The functionally-annotated GENCODE v39 reference transcriptome available at the tappAS website, was supplied to the --gff3 flag. A tappAS-compatible GFF3 file was obtained as a result. In addition, taking advantage of the Kallisto quantification carried out internally by SQANTI3 QC, an expression matrix was created using the TPM values of each transcript model calculated per replicate. Finally, an experimental design table was generated specifying the conditions (i.e. differentiation stage)of each replicate. These three files (functional annotation GFF3 file, expression, and design matrices) were used to create a project in tappAS and perform Differential Expression, Differential Isoform Usage, and Functional Enrichment analyses.

#### 4.3.6 SIRV evaluation metrics

To assess the ability of SQANTI3 to accept or reject transcripts accurately, Lexogen SIRVs introduced during library preparation were used. Since SQANTI3 uses the reference annotation to categorize the transcript models and this affects the curation process, the annotation of the SIRVs was modified to have ”novel” isoforms and isoforms annotated but not present in the sample. To achieve it, the annotation of the SIRV genes was added to the GENCODE human reference transcriptome (v39), including the insufficient and the over-annotated versions available at the Lexogen website https://www.lexogen.com/wp-content/uploads/2021/06/SIRV_Set4_Norm_Sequences_20210507.zip. We considered an isoform as True Positive (TP) if it matched a transcript model of the complete and correct annotation as an FSM Reference Match. Depending on the spike-in matched, it could be a known or novel TP if it was present in the modified annotation or not, respectively. When the transcript model could be associated with the transcript present in the sample but differed by more than 50bp in any of their ends, this was counted as a Partial TP (PTP). Transcripts matching a false SIRV present in the reference were computed as an over-annotation FP. The rest of the transcripts that were classified as NIC or NNC compared to any annotation (complete or modified) were reported as False Positives (FP). Moreover, we measured the number of isoforms considered novel during the SQANTI3 curation process which includes novel TP and FP. Taking these figures, sensitivity, precision, F-score, False Discovery Rate (FDR), Over-annotation Detection Rate (ODR), and Novel

Detection Rate (NDR) were calculated with the following formulas:

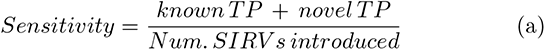

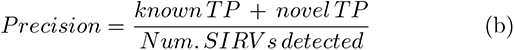

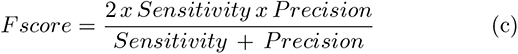

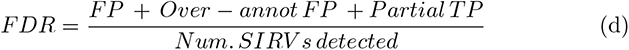

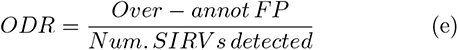

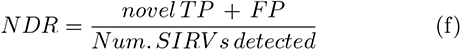

## 5 Data and Code availability

The SQANTI3 software is available at https://github.com/ConesaLab/SQANTI3. All the files used to generate the results in this paper are publicly accessible at http://conesalab.org/SQANTI3/. For an easier exploration of WTC11 transcript models identified with IsoSeq3 and their characterization with SQANTI3, a specific and public Track hub was generated for the UCSC Genome Browser (hub URL: http://conesalab.org/SQANTI3/WTC11/SQANTI3_hub/hub.txt), including the orthogonal data used for validation. We make a special emphasis on the availability of the results of hESC H1 cells to endoderm differentiation in a ready-to-use format on tappAS (http://conesalab.org/SQANTI3/H1endo/tappAS_files/).

## Acknowledgments

This work has been funded by NIH grant R21HG011280, by the Spanish Ministry of Science grants BES-2016-076994 and PID2020-119537RB-100 and by the Comunitat Valenciana grant ACIF/2018/290.

## 6 Author Contributions

F.P.P. developed and implemented novel SQANTI3 QC features, developed and expanded the SQANTI3 rules filter, designed and performed data analyses, and drafted the manuscript. A.A.L. refactored and improved the SQANTI3 ML filter, developed and implemented the SQANTI3 rescue module, and drafted the manuscript. L.M. contributed to SQANTI3 QC implementations. P.S. developed and implemented IsoAnnotLite, J.M.L contributed to data analysis, R.A. contributed to novel SQANTI3 QC graphical output, E.E.M. contributed to SQANTI3 ML filter development, T.L. contributed to SQANTI3 QC graphical output, A.N. contributed to Rescue validation analyses, L.McI. contributed to conceptualization, E.T. contributed to defining SQANTI3 categories, developed usage of additional data in SQANTI3 QC, and conceived the rules filter strategies. A.C. conceived and supervised the study. All authors contributed to drafting the manuscript.

## Appendix A Supplementary Figures

**Fig. A1.**
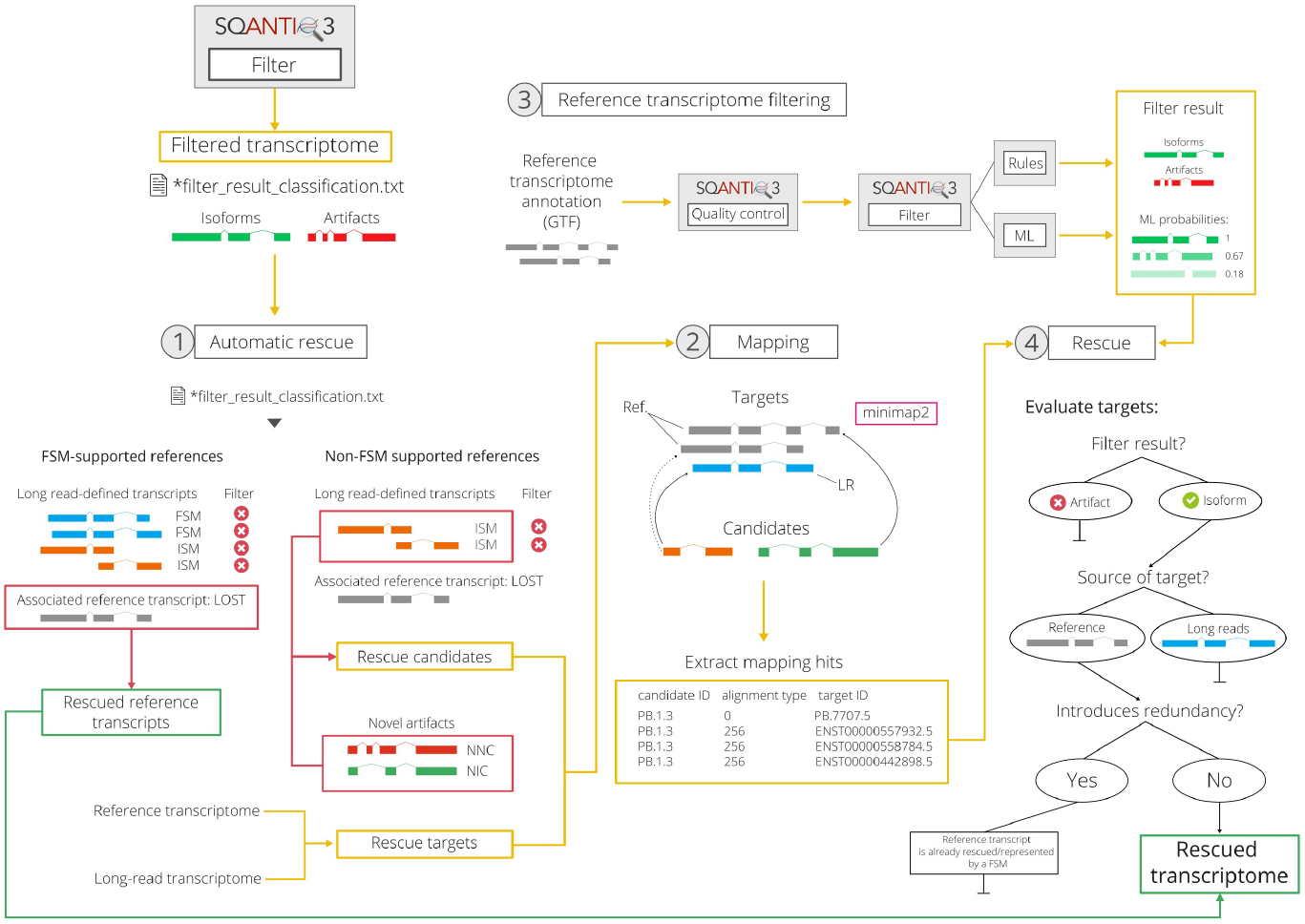
Supplementary Figure 1. SQANTI3 Rescue workflow. 1) If an FSM-supported reference transcript is lost during the filtering, the version of the reference is automatically rescued. 2) The rest of the LR-defined transcript models filtered out (rescue candidates) are mapped against the reference transcriptome combined with the accepted LR-defined isoforms (rescue targets), allowing several hits per candidate. 3) Reference transcriptome was previously evaluated and filtered with the same data and criteria as the LR-defined transcripts. 4) Rescue is completed by evaluating targets. They need to pass the filtering and do not increase the redundancy, meaning that if the target is an LR-defined transcript present or it is a reference transcript already represented as an FSM in the filtered transcriptome, these targets will not be added to the final annotation.

**Fig. A2.**
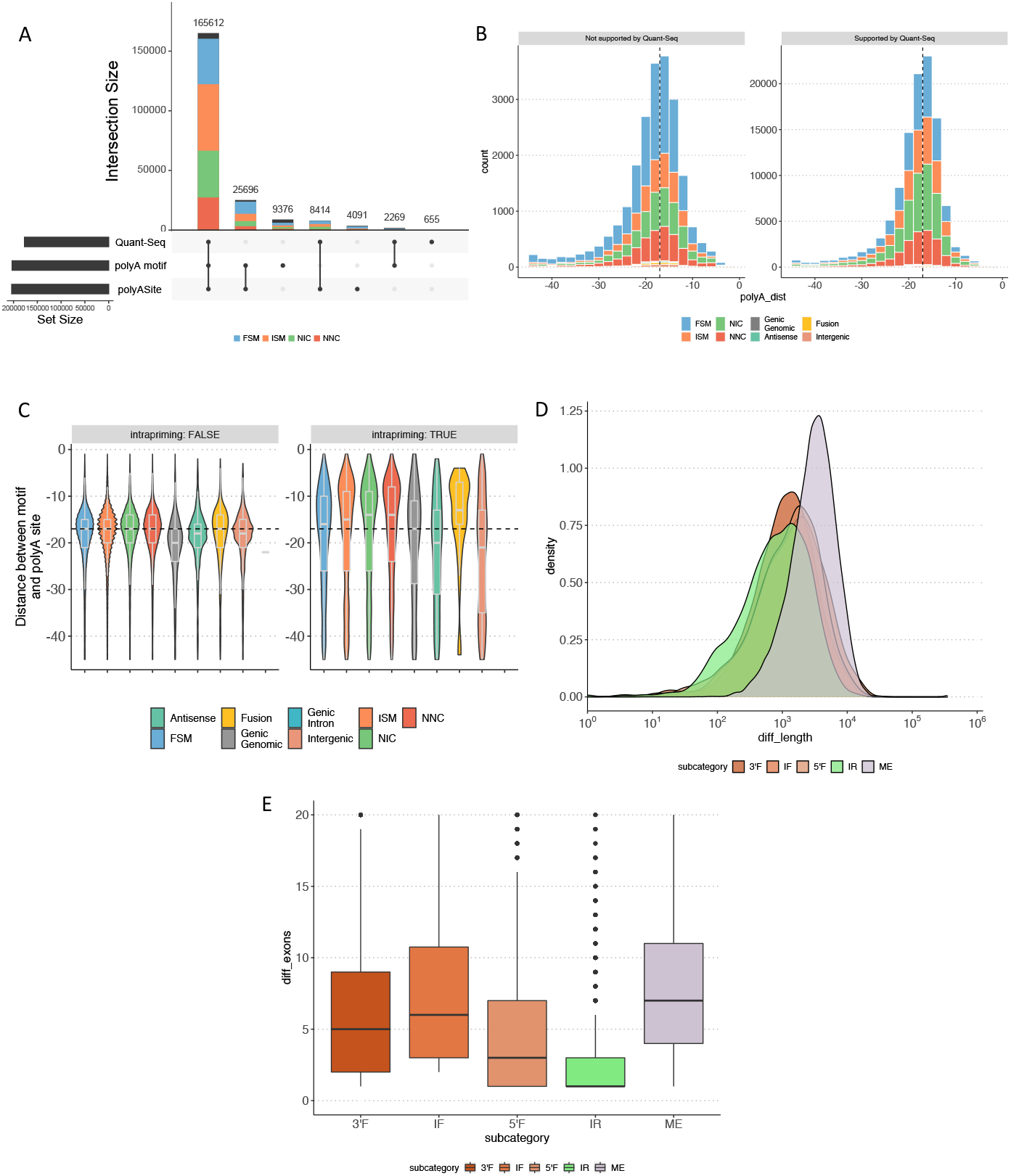
Supplementary Figure 2. A) Agreement validating detected TTSs using different sources of additional information: sample-specific Quant-Seq data, presence of polyA motif and polyASite database. B) Number of isoforms (y-axis), stratified by structural category, compared against the distance between their polyA motifs and polyA sites (x-axis). Isoforms were divided depending on their support by Quant-Seq data. B) Distribution of distances between polyA motifs and polyA sites, stratified by structural categories. Isoforms were divided depending if they could be considered intrapriming artifacts. D) Distribution of length differences between ISM isoforms (stratified by subcategory) and their reference counterparts. E) Distribution of differences of exon number between ISM isoforms (stratified by subcategory) and their reference counterparts.

**Fig. A3.**
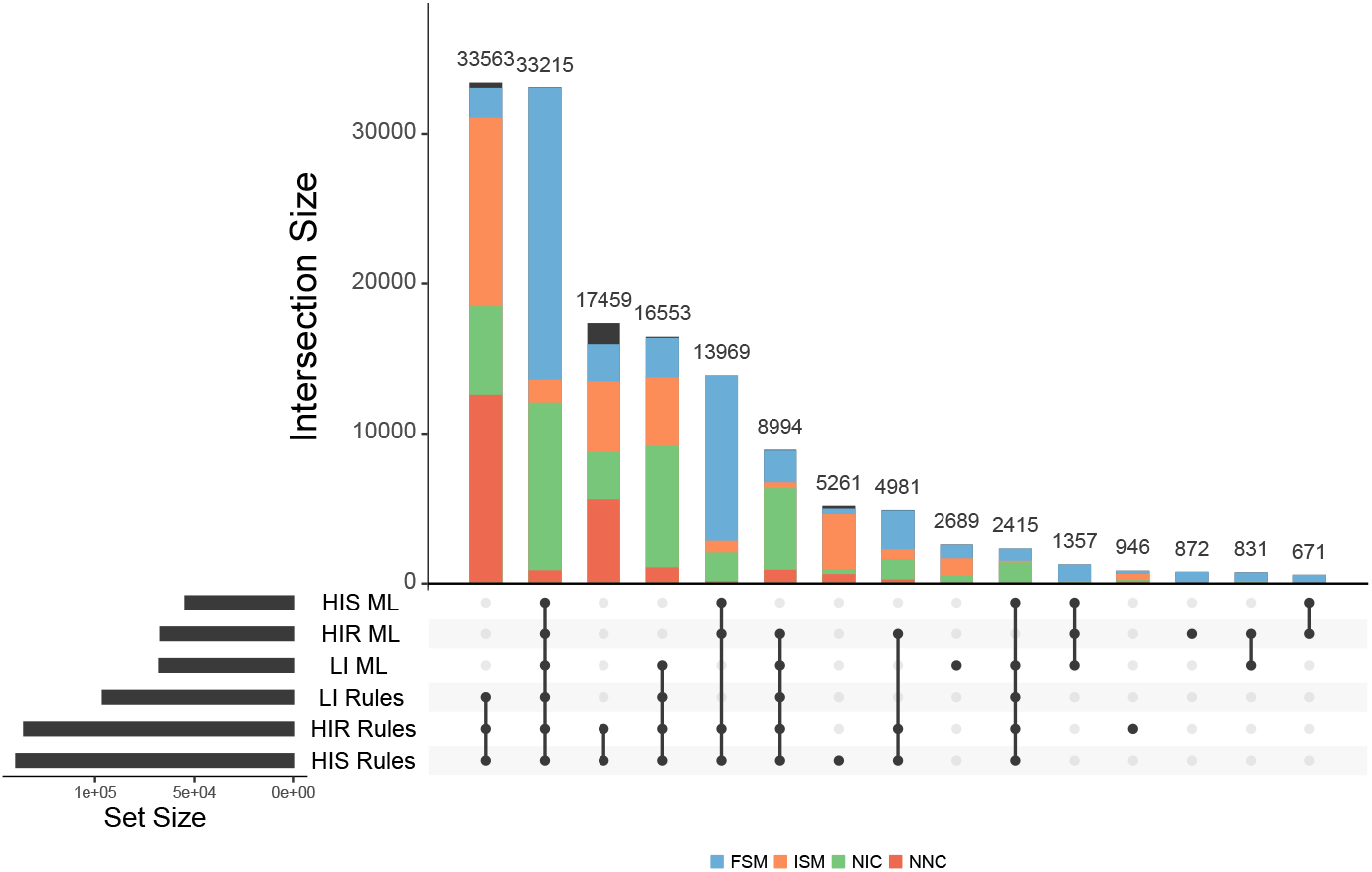
Extended Data Figure 1. Agreement between filtered transcriptomes relied on the strategy and data used.

**Fig. A4.**
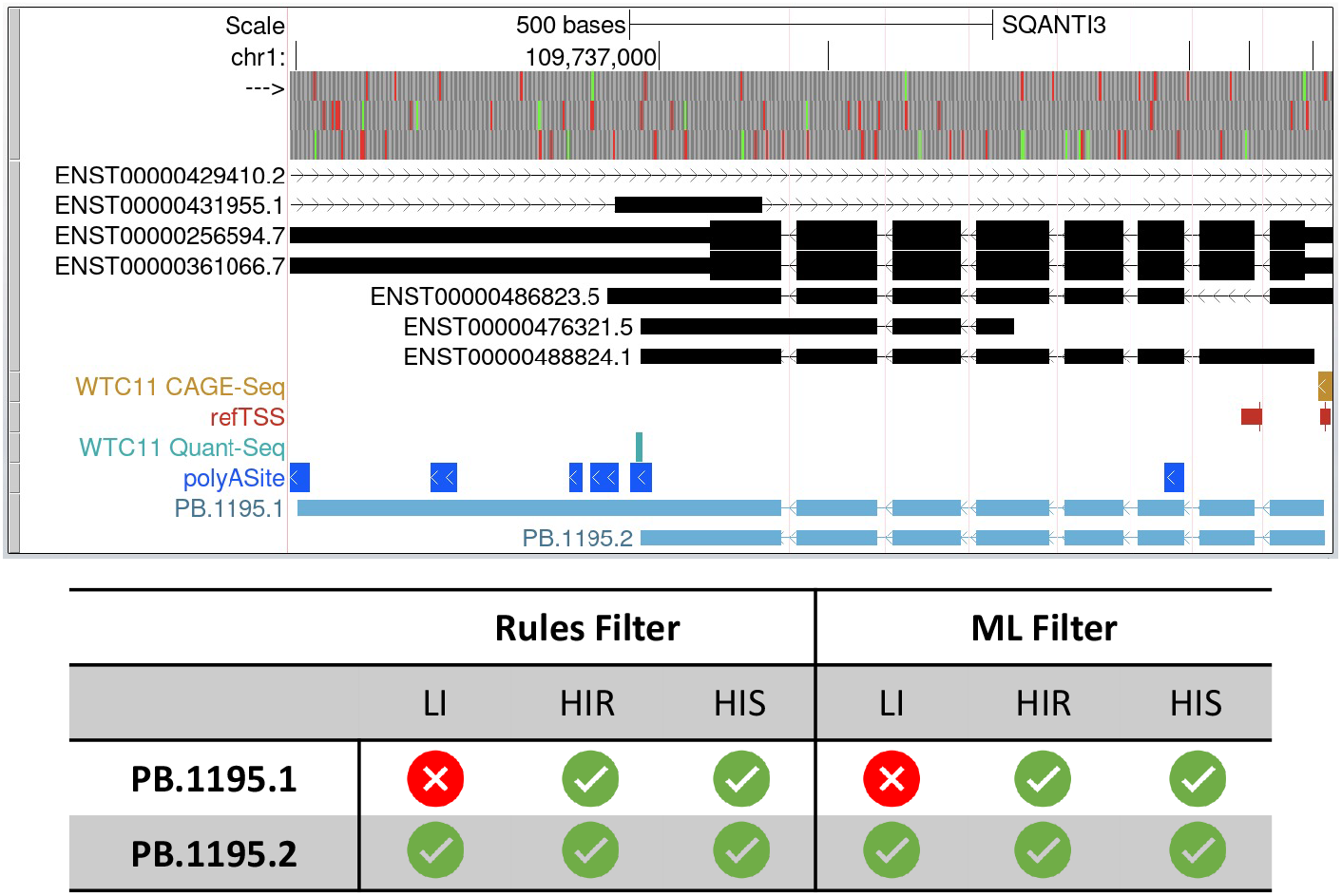
Supplementary Figure 3. UCSC Genome Browser view of human GST5 gene. Upper track (black) represents the GENCODE annotation for that locus, which includes 5 isoforms for the GST5 gene coded in the negative strand. The bottom track (light blue) shows the LR-defined isoforms identified in the WTC11 sample, while the tracks in between (gold, red, turquoise, and dark blue) are the orthogonal data available to validate those isoforms. The bottom table indicates which filters passed each of the isoforms regarding the informative situation simulated: Low Input (LI), High Input Reference (HIR), and High Input Sample-specific (HIS).

**Fig. A5.**
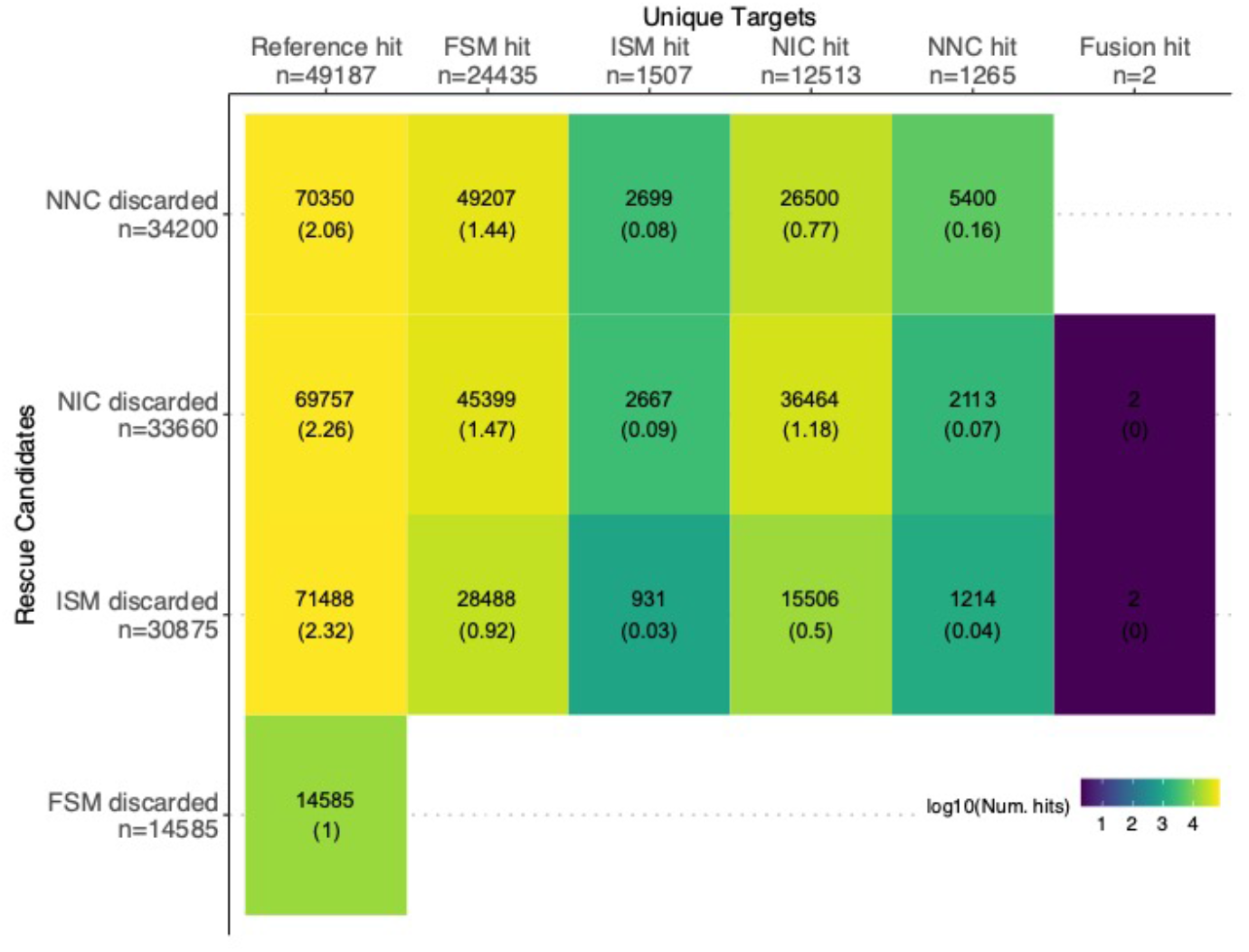
Supplementary Figure 4. Relationship between discarded transcripts (rescue candidates) and their rescue targets in the ML-HIS filtering scenario.

**Fig. A6.**
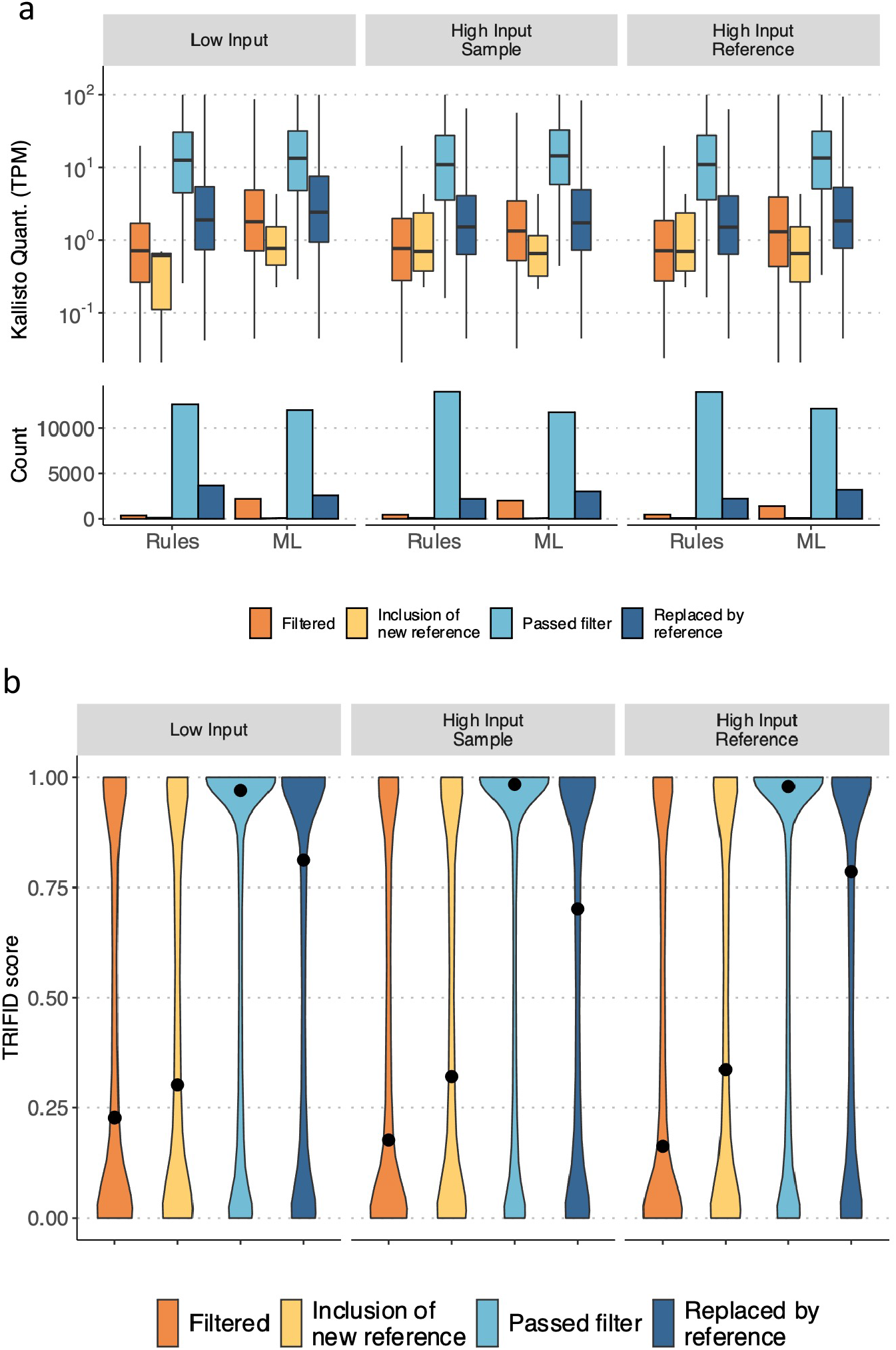
Extended Data Figure 2. Distribution of **a)** expression values (TPM) of known genes and **b)** TRIFID scores of known transcripts identified. Filtered genes/transcripts (orange) did not pass the corresponding filter and were not eventually rescued. Genes/- transcripts filtered but recovered by introducing an isoform from the reference (dark blue) represent the rescue strategy’s fundamental purpose. In exceptional cases, genes/transcripts models not initially detected were included in the final transcriptome (yellow) via rescue.

**Fig. A7.**
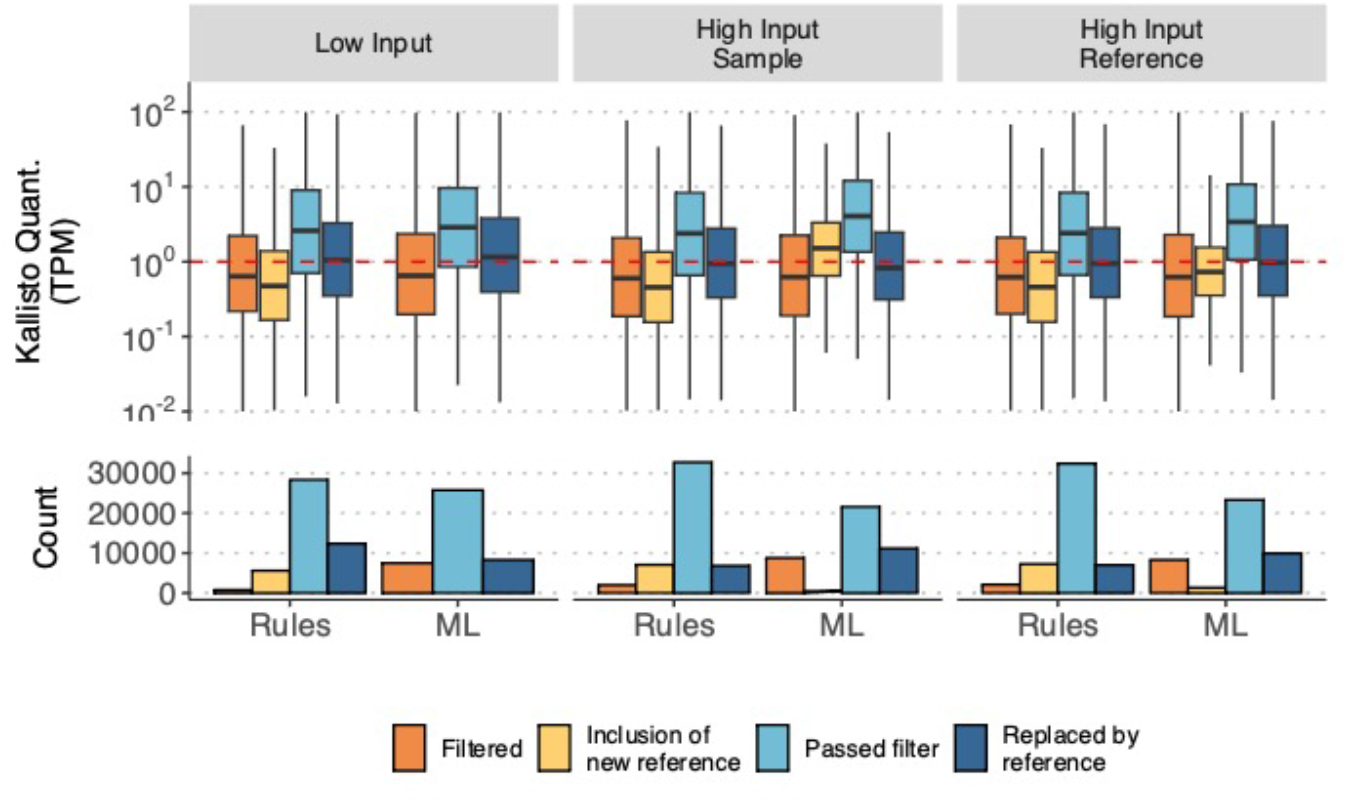
Supplementary Figure 5. Distribution of expression values (TPM) of known transcripts -detected as FSM or ISM- for each filtering and rescue scenario. Filtered isoforms (orange) were those FSM/ISM that did not pass the corresponding filter and were not eventually rescued because of unacceptable quality attributes. On the other hand, LR-defined known transcripts lost during filtering but recovered by introducing transcript models from the reference (dark blue) are the the rescue strategy’s goal. In some exceptional cases, genes not initially detected were included in the final transcriptome (yellow) if a discarded sequence mapped them and the filtering criteria were fulfilled.

**Fig. A8.**
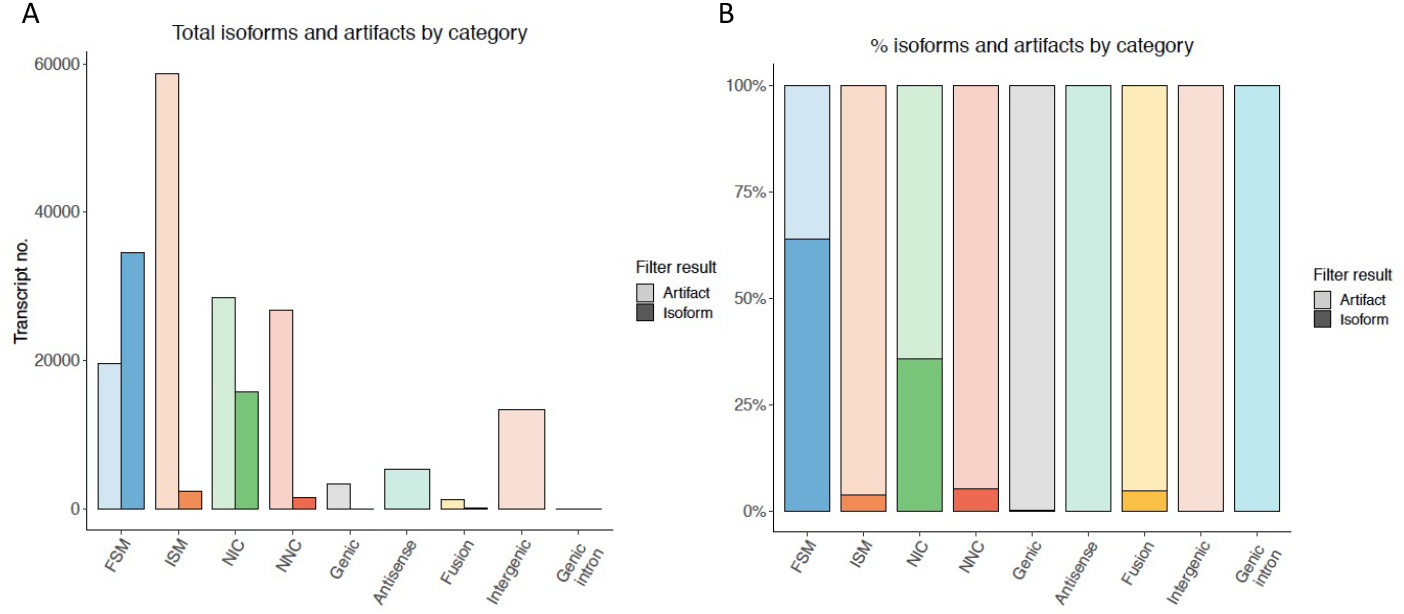
Supplementary Figure 5. Distribution of isoforms and artifacts by structural category identified in the hESC H1-endoderm differentiation experiment, after running a ML-based filter. A) Absolute number of isoforms classified as isoforms or artifacts per structural category. B) Percentage of isoforms and artifacts within each structural category.

**Fig. A9.**
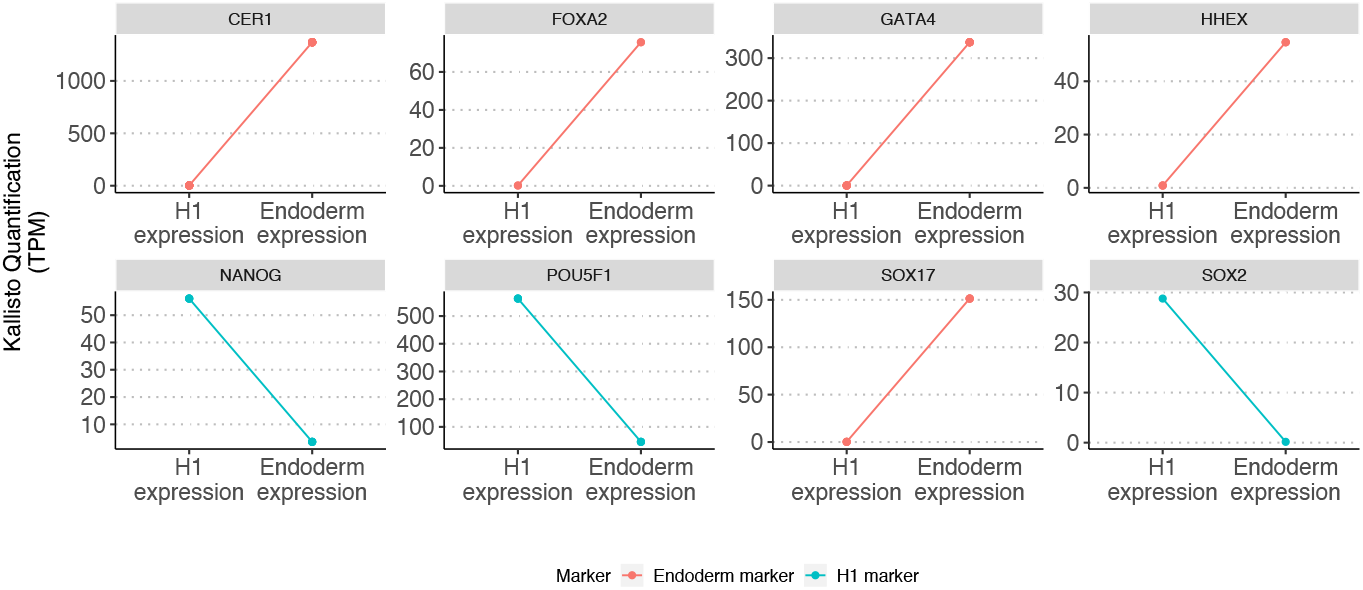
Supplementary Figure 6. Gene expression (TPM) of hESC H1-endoderm differentiation markers calculated using Kallisto.

**Fig. A10.**
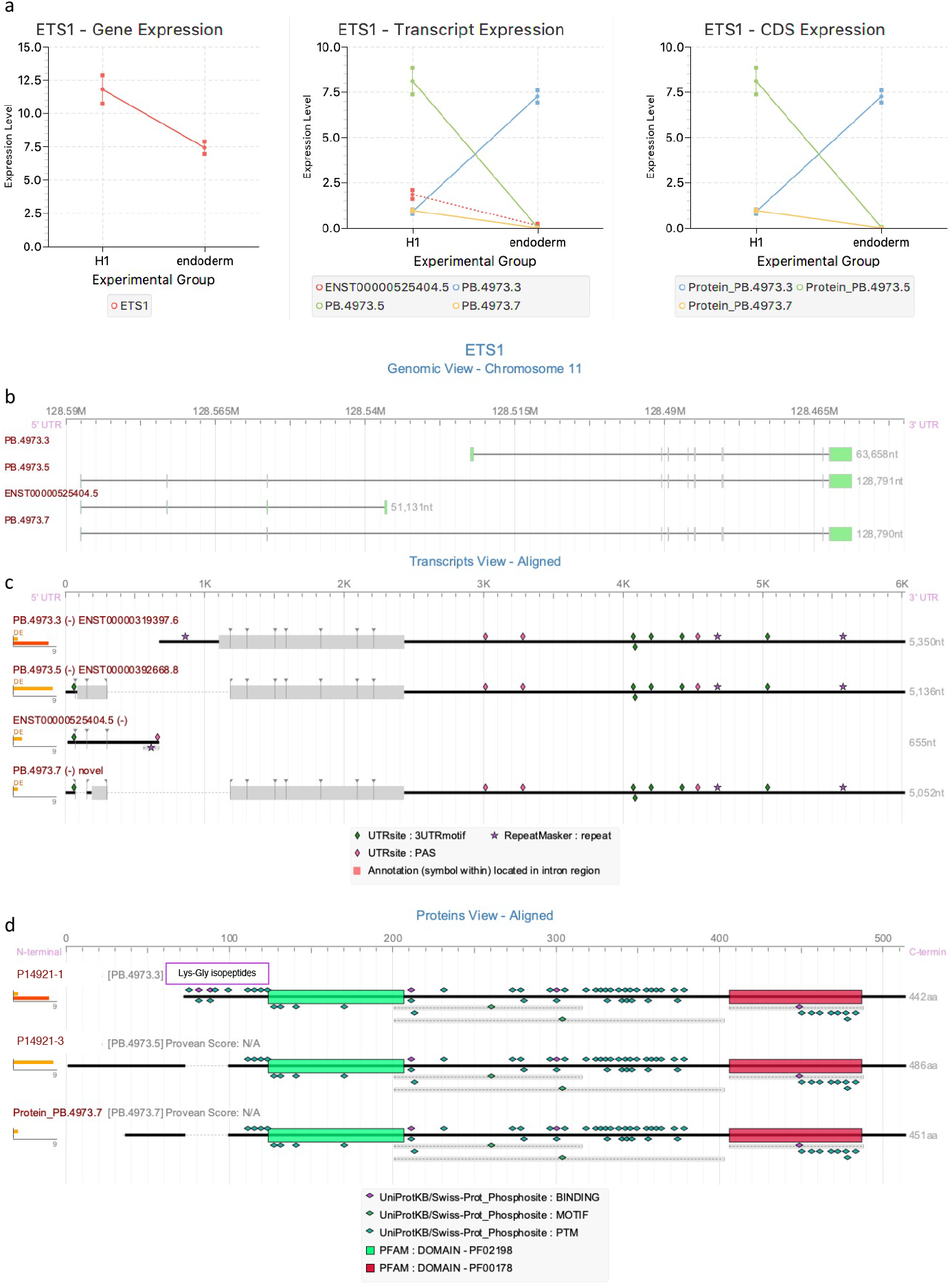
Extended Data Figure 3. tappAS visualization of ETS1 gene through hESC H1- endoderm differentiation. A) Expression values (TPM) at the gene, transcript and CDS levels. Dotted line in transcript expression plot represent a non-coding isoform. B) Genomic View. Green segments represent untranslated regions. C) Transcript View. Thick sections represent the location of predicted ORFs. D) Protein View.

**Fig. A11.**
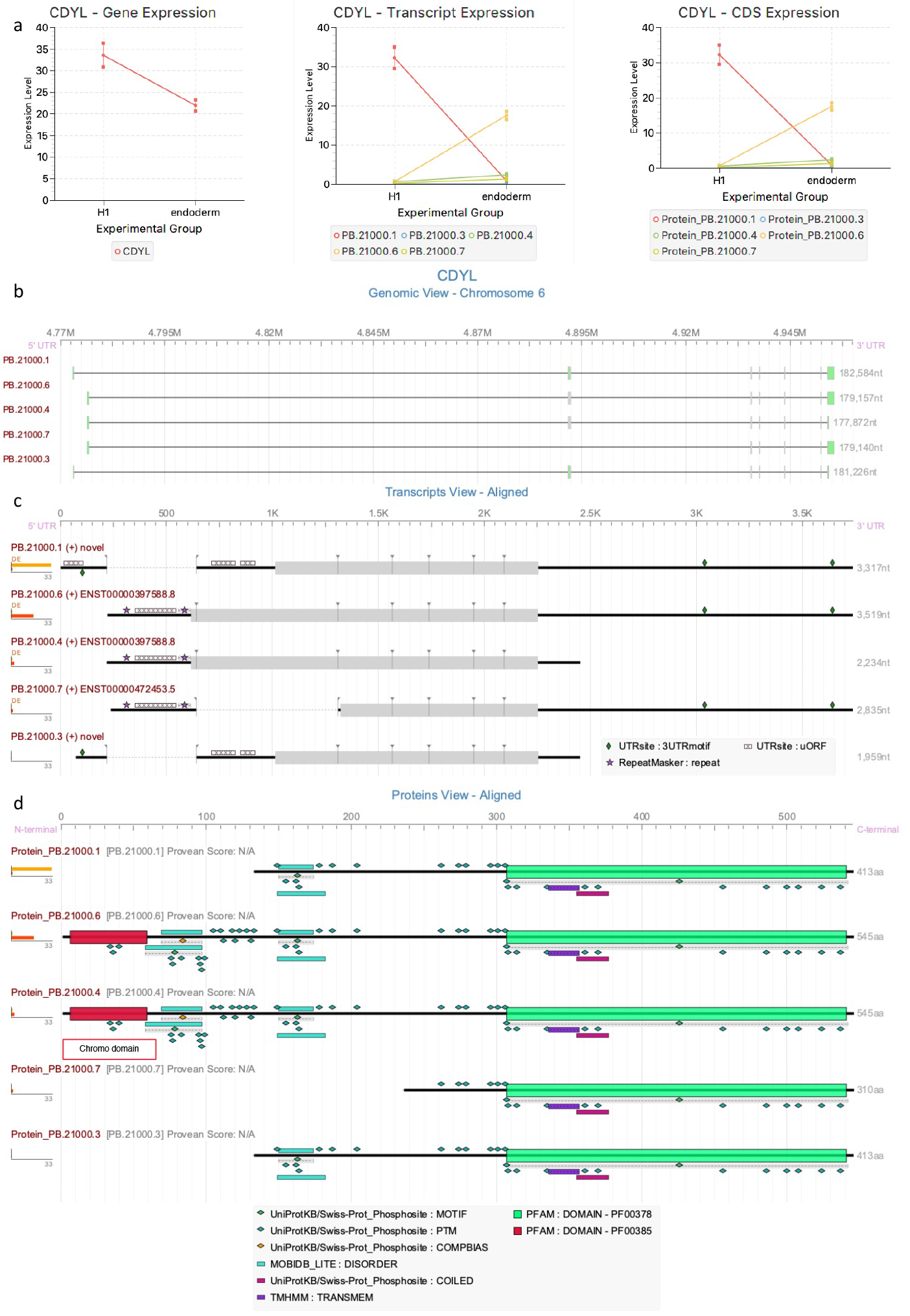
Extended Data Figure 4. tappAS visualization of CDYL gene through hESC H1-endoderm differentiation. A) Expression values (TPM) at the gene, transcript and CDS levels. B) Genomic View. Green segments represent untranslated regions. C) Transcript View. Thick sections represent the location of predicted ORFs. D) Protein View.

**Table A1.**
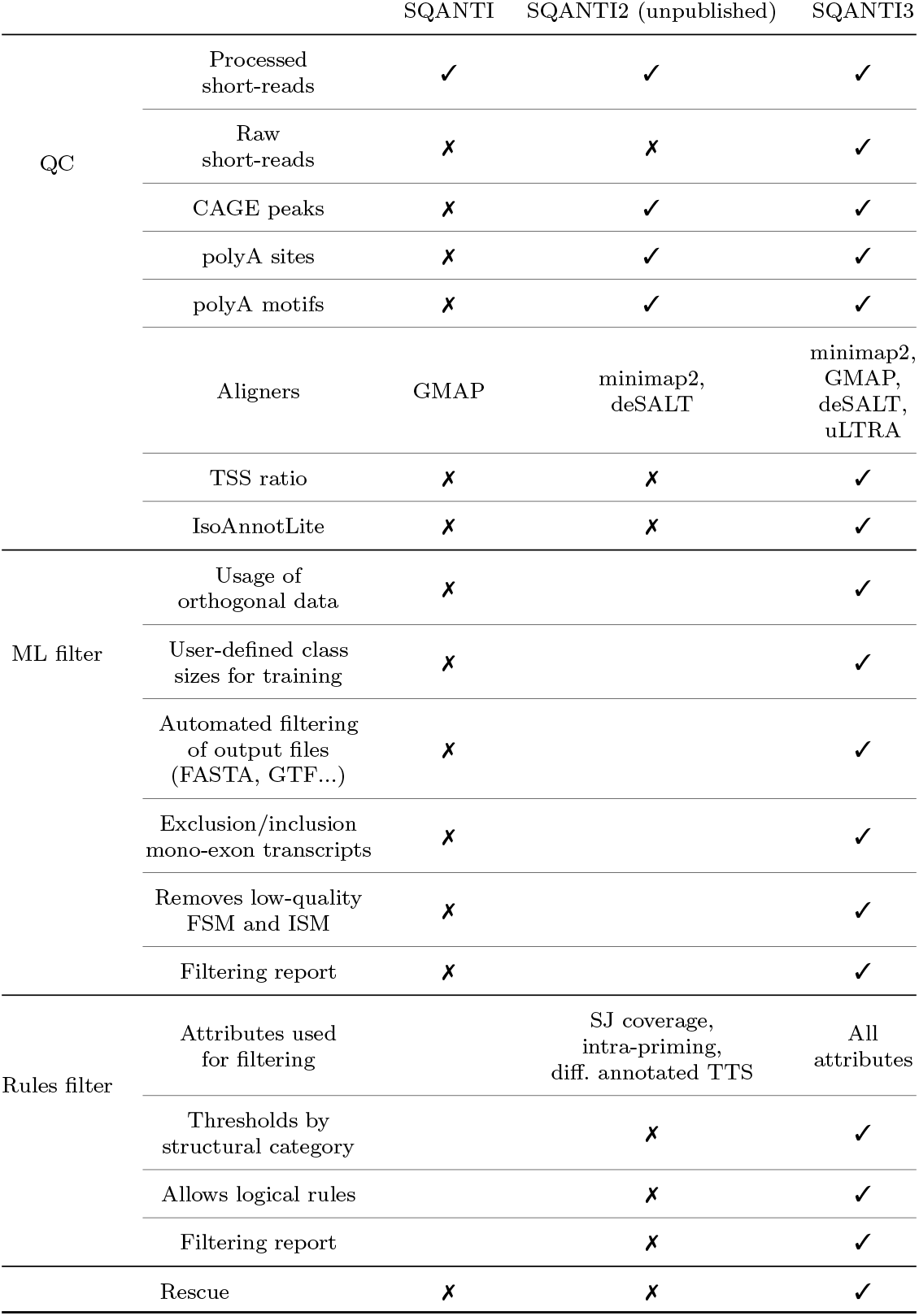
Supplementary Table 1. Differences and upgrades of SQANTI throughout all its versions since its first release until the current publication

## References

[1] Method of the year 2022: long-read sequencing 20(1), 1–1. https://doi.org/10.1038/s41592-022-01759-x. Number: 1 Publisher: Nature Publishing Group. Accessed 2023-02-06

[2] Ding, C., Yan, X., Xu, M., Zhou, R., Zhao, Y., Zhang, D., Huang, Z., Pan, Z., Xiao, P., Li, H., Chen, L., Wang, Y.: Short-read and long-read full-length transcriptome of mouse neural stem cells across neurodevelopmental stages 9(1), 69. https://doi.org/10.1038/s41597-022-01165-0. Number: 1 Publisher: Nature Publishing Group. Accessed 2022-11-08

[3] Tilgner, H., Grubert, F., Sharon, D., Snyder, M.P.: Defining a personal, allele-specific, and single-molecule long-read transcriptome 111(27), 9869–9874. https://doi.org/10.1073/pnas.1400447111. Publisher: Proceedings of the National Academy of Sciences. Accessed 2022-11-08

[4] Singh, M., Al-Eryani, G., Carswell, S., Ferguson, J.M., Blackburn, J., Barton, K., Roden, D., Luciani, F., Giang Phan, T., Junankar, S., Jackson, K., Goodnow, C.C., Smith, M.A., Swarbrick, A.: High-throughput targeted long-read single cell sequencing reveals the clonal and transcriptional landscape of lymphocytes 10(1), 3120. https://doi.org/10.1038/s41467-019-11049-4. Number: 1 Publisher: Nature Publishing Group. Accessed 2022-11-08

[5] Wang, B., Tseng, E., Regulski, M., Clark, T.A., Hon, T., Jiao, Y., Lu, Z., Olson, A., Stein, J.C., Ware, D.: Unveiling the complexity of the maize transcriptome by single-molecule long-read sequencing 7(1), 11708. https://doi.org/10.1038/ncomms11708. Number: 1 Publisher: Nature Publishing Group. Accessed 2022-11-08

[6] Hasan, S., Huang, L., Liu, Q., Perlo, V., O’Keeffe, A., Margarido, G.R.A., Furtado, A., Henry, R.J.: The long read transcriptome of rice (oryza sativa ssp. japonica var. nipponbare) reveals novel transcripts 15(1), 29. https://doi.org/10.1186/s12284-022-00577-1. Accessed 2022-11-08

[7] Wang, B., Tseng, E., Baybayan, P., Eng, K., Regulski, M., Jiao, Y., Wang, L., Olson, A., Chougule, K., Buren, P.V., Ware, D.: Variant phasing and haplotypic expression from long-read sequencing in maize 3(1), 1–11. https://doi.org/10.1038/s42003-020-0805-8. Number: 1 Publisher: Nature Publishing Group. Accessed 2022-11-08

[8] Wang, Y., Wang, H., Xi, F., Wang, H., Han, X., Wei, W., Zhang, H., Zhang, Q., Zheng, Y., Zhu, Q., Kohnen, M.V., Reddy, A.S.N., Gu, L.: Profiling of circular RNA n6-methyladenosine in moso bamboo (phyllostachys edulis) using nanopore-based direct RNA sequencing 62(12), 1823–1838. https://doi.org/10.1111/jipb.13002. eprint: https://onlinelibrary.wiley.com/doi/pdf/10.1111/jipb.13002. Accessed 2022-11-08

[9] Gupta, I., Collier, P.G., Haase, B., Mahfouz, A., Joglekar, A., Floyd, T., Koopmans, F., Barres, B., Smit, A.B., Sloan, S.A., Luo, W., Fedrigo, O., Ross, M.E., Tilgner, H.U.: Single-cell isoform RNA sequencing characterizes isoforms in thousands of cerebellar cells. https://doi.org/10.1038/nbt.4259

[10] Dai, Z., Ren, J., Tong, X., Hu, H., Lu, K., Dai, F., Han, M.-J.: The landscapes of full-length transcripts and splice isoforms as well as transposons exonization in the lepidopteran model system, bombyx mori 12, 704162. https://doi.org/10.3389/fgene.2021.704162

[11] Lu, P., Chen, D., Qi, Z., Wang, H., Chen, Y., Wang, Q., Jiang, C., Xu, J.-R., Liu, H.: Landscape and regulation of alternative splicing and alternative polyadenylation in a plant pathogenic fungus 235(2), 674–689. https://doi.org/10.1111/nph.18164

[12] Gao, C., Ren, L., Wang, M., Wang, Z., Fu, N., Wang, H., Shi, J.: Fulllength transcriptome sequencing-based analysis of pinus sylvestris var. mongolica in response to sirex noctilio venom 13(4), 338. https://doi.org/10.3390/insects13040338

[13] Wyman, D., Balderrama-Gutierrez, G., Reese, F., Jiang, S., Rahmanian, S., Forner, S., Matheos, D., Zeng, W., Williams, B., Trout, D., England, W., Chu, S.-H., Spitale, R.C., Tenner, A.J., Wold, B.J., Mortazavi, A.: A technology-agnostic long-read analysis pipeline for transcriptome discovery and quantification. bioRxiv. Pages: 672931 Section: New Results. https://doi.org/10.1101/672931. https://www.biorxiv.org/content/10.1101/672931v2 Accessed 2022-11-15

[14] Kovaka, S., Zimin, A.V., Pertea, G.M., Razaghi, R., Salzberg, S.L., Pertea, M.: Transcriptome assembly from long-read RNA-seq alignments with StringTie2 20(1), 278. https://doi.org/10.1186/s13059-019-1910-1. Accessed 2022-11-15

[15] Volden, R., Palmer, T., Byrne, A., Cole, C., Schmitz, R.J., Green, R.E., Vollmers, C.: Improving nanopore read accuracy with the r2c2 method enables the sequencing of highly multiplexed full-length single-cell cDNA 115(39), 9726–9731. https://doi.org/10.1073/pnas.1806447115

[16] Tian, L., Jabbari, J.S., Thijssen, R., Gouil, Q., Amarasinghe, S.L., Voogd, O., Kariyawasam, H., Du, M.R.M., Schuster, J., Wang, C., Su, S., Dong, X., Law, C.W., Lucattini, A., Prawer, Y.D.J., Collar-Fernández, C., Chung, J.D., Naim, T., Chan, A., Ly, C.H., Lynch, G.S., Ryall, J.G., Anttila, C.J.A., Peng, H., Anderson, M.A., Flensburg, C., Majewski, I., Roberts, A.W., Huang, D.C.S., Clark, M.B., Ritchie, M.E.: Comprehensive characterization of single-cell full-length isoforms in human and mouse with long-read sequencing 22(1), 310. https://doi.org/10.1186/s13059-021-02525-6. Accessed 2022-11-15

[17] Prjibelski, A., Mikheenko, A., Joglekar, A., Smetanin, A., Jarroux, J., Lapidus, A., Tilgner, H.: IsoQuant: a Tool for Accurate Novel Isoform Discovery with Long Reads. https://doi.org/10.21203/rs.3.rs-1571850/v1. https://www.researchsquare.com/article/rs-1571850/v1 Accessed 2022-11-15

[18] Tang, A.D., Soulette, C.M., van Baren, M.J., Hart, K., Hrabeta-Robinson, E., Wu, C.J., Brooks, A.N.: Full-length transcript characterization of SF3b1 mutation in chronic lymphocytic leukemia reveals downregulation of retained introns 11(1), 1438. https://doi.org/10.1038/s41467-020-15171-6. Number: 1 Publisher: Nature Publishing Group. Accessed 2022-11-15

[19] de la Fuente, L., Arzalluz-Luque,, Tardáguila, M., del Risco, H., Martí, C., Tarazona, S., Salguero, P., Scott, R., Lerma, A., Alastrue-Agudo, A., Bonilla, P., Newman, J.R.B., Kosugi, S., McIntyre, L.M., Moreno-Manzano, V., Conesa, A.: tappAS: a comprehensive computational framework for the analysis of the functional impact of differential splicing 21(1), 119. https://doi.org/10.1186/s13059-020-02028-w. Accessed 2022-11-08

[20] Weirather, J.L., Cesare, M.d., Wang, Y., Piazza, P., Sebastiano, V., Wang, X.-J., Buck, D., Au, K.F.: Comprehensive Comparison of Pacific Biosciences and Oxford Nanopore Technologies and Their Applications to Transcriptome Analysis. https://doi.org/10.12688/f1000research.10571.2. Type: article. https://f1000research.com/articles/6-100 Accessed 2022-12-20

[21] Wang, Y., Zhao, Y., Bollas, A., Wang, Y., Au, K.F.: Nanopore sequencing technology, bioinformatics and applications 39(11), 1348–1365. https://doi.org/10.1038/s41587-021-01108-x. Number: 11 Publisher: Nature Publishing Group. Accessed 2023-01-19

[22] Hon, T., Mars, K., Young, G., Tsai, Y.-C., Karalius, J.W., Landolin, J.M., Maurer, N., Kudrna, D., Hardigan, M.A., Steiner, C.C., Knapp, S.J., Ware, D., Shapiro, B., Peluso, P., Rank, D.R.: Highly accurate long-read HiFi sequencing data for five complex genomes 7(1), 399. https://doi.org/10.1038/s41597-020-00743-4. Number: 1 Publisher: Nature Publishing Group. Accessed 2023-02-06

[23] Tardaguila, M., de la Fuente, L., Marti, C., Pereira, C., Pardo-Palacios, F.J., del Risco, H., Ferrell, M., Mellado, M., Macchietto, M., Verheggen, K., Edelmann, M., Ezkurdia, I., Vazquez, J., Tress, M., Mortazavi, A., Martens, L., Rodriguez-Navarro, S., Moreno-Manzano, V., Conesa, A.: SQANTI: extensive characterization of long-read transcript sequences for quality control in full-length transcriptome identification and quantification 28(3), 396–411. https://doi.org/10.1101/gr.222976.117. Accessed 2022-10-22

[24] Ray, T.A., Cochran, K., Kozlowski, C., Wang, J., Alexander, G., Cady, M.A., Spencer, W.J., Ruzycki, P.A., Clark, B.S., Laeremans, A., He, M.-X., Wang, X., Park, E., Hao, Y., Iannaccone, A., Hu, G., Fedrigo, O., Skiba, N.P., Arshavsky, V.Y., Kay, J.N.: Comprehensive identification of mRNA isoforms reveals the diversity of neural cell-surface molecules with roles in retinal development and disease 11(1), 3328. https://doi.org/10.1038/s41467-020-17009-7. Number: 1 Publisher: Nature Publishing Group. Accessed 2022-12-21

[25] Palmer, C.R., Liu, C.S., Romanow, W.J., Lee, M.-H., Chun, J.: Altered cell and RNA isoform diversity in aging down syndrome brains 118(47), 2114326118. https://doi.org/10.1073/pnas.2114326118. Publisher: Proceedings of the National Academy of Sciences. Accessed 2022-12-21

[26] Miller, R.M., Jordan, B.T., Mehlferber, M.M., Jeffery, E.D., Chatzipantsiou, C., Kaur, S., Millikin, R.J., Dai, Y., Tiberi, S., Castaldi, P.J., Shortreed, M.R., Luckey, C.J., Conesa, A., Smith, L.M., Deslattes Mays, A., Sheynkman, G.M.: Enhanced protein isoform characterization through long-read proteogenomics 23(1), 69. https://doi.org/10.1186/s13059-022-02624-y. Accessed 2022-12-21

[27] Tseng, E., Underwood, J.G., Evans Hutzenbiler, B.D., Trojahn, S., Kingham, B., Shevchenko, O., Bernberg, E., Vierra, M., Robbins, C.T., Jansen, H.T., Kelley, J.L.: Long-read isoform sequencing reveals tissue-specific isoform expression between active and hibernating brown bears (ursus arctos) 12(3), 422. https://doi.org/10.1093/g3journal/jkab422. Accessed 2022-12-21

[28] Takahashi, H., Lassmann, T., Murata, M., Carninci, P.: 5’ end-centered expression profiling using cap-analysis gene expression and next-generation sequencing 7(3), 542–561. https://doi.org/10.1038/nprot.2012.005

[29] Moll, P., Ante, M., Seitz, A., Reda, T.: QuantSeq 3 mRNA sequencing for RNA quantification 11(12). https://doi.org/10.1038/nmeth.f.376. Number: 12 Publisher: Nature Publishing Group. Accessed 2022-12-21

[30] Veeneman, B.A., Shukla, S., Dhanasekaran, S.M., Chinnaiyan, A.M., Nesvizhskii, A.I.: Two-pass alignment improves novel splice junction quantification 32(1), 43–49. https://doi.org/10.1093/bioinformatics/btv642. Accessed 2022-12-10

[31] Pardo-Palacios, F., Reese, F., Carbonell-Sala, S., Diekhans, M., Liang, C., Wang, D., Williams, B., Adams, M., Behera, A., Lagarde, J., Li, H., Prjibelski, A., Balderrama-Gutierrez, G., Çelik, M.H., De María, M., Denslow, N., Garcia-Reyero, N., Goetz, S., Hunter, M., Loveland, J., Menor, C., Moraga, D., Mudge, J., Takahashi, H., Tang, A., Youngworth, I., Carninci, P., Guigó, R., Tilgner, H., Wold, B., Vollmers, C., Sheynkman, G., Frankish, A., Au, K.F., Conesa, A., Mortazavi, A., Brooks, A.N.: Systematic Assessment of Long-read RNA-seq Methods for Transcript Identification and Quantification. https://doi.org/10.21203/rs.3.rs-777702/v1. https://www.researchsquare.com/article/rs-777702/v1 Accessed 2022-12-21

[32] Abugessaisa, I., Noguchi, S., Hasegawa, A., Kondo, A., Kawaji, H., Carninci, P., Kasukawa, T.: refTSS: A reference data set for human and mouse transcription start sites 431(13), 2407–2422. https://doi.org/10.1016/j.jmb.2019.04.045. Accessed 2022-12-21

[33] Beaudoing, E., Freier, S., Wyatt, J.R., Claverie, J.M., Gautheret, D.: Patterns of variant polyadenylation signal usage in human genes 10(7), 1001–1010. https://doi.org/10.1101/gr.10.7.1001

[34] Rodriguez, J.M., Pozo, F., Cerdán-Vélez, D., Di Domenico, T., Vázquez, J., Tress, M.: APPRIS: selecting functionally important isoforms 50, 54–59. https://doi.org/10.1093/nar/gkab1058. Accessed 2022-12-29

[35] Paul, L., Kubala, P., Horner, G., Ante, M., Holländer, I., Alexander, S., Reda, T.: SIRVs: Spike-In RNA Variants as External Isoform Controls in RNA-Sequencing. bioRxiv. Pages: 080747 Section: New Results. https://doi.org/10.1101/080747. https://www.biorxiv.org/content/10.1101/080747v1 Accessed 2022-12-29

[36] Grillo, G., Turi, A., Licciulli, F., Mignone, F., Liuni, S., Banfi, S., Gennarino, V.A., Horner, D.S., Pavesi, G., Picardi, E., Pesole, G.: UTRdb and UTRsite (RELEASE 2010): a collection of sequences and regulatory motifs of the untranslated regions of eukaryotic mRNAs 38, 75–80. https://doi.org/10.1093/nar/gkp902. Accessed 2022-12-07

[37] Siller, R., Naumovska, E., Mathapati, S., Lycke, M., Greenhough, S., Sullivan, G.J.: Development of a rapid screen for the endodermal differentiation potential of human pluripotent stem cell lines 6(1), 37178. https://doi.org/10.1038/srep37178. Number: 1 Publisher: Nature Publishing Group. Accessed 2023-01-05

[38] Baumgart, E., Vanhooren, J.C., Fransen, M., Marynen, P., Puype, M., Vandekerckhove, J., Leunissen, J.A., Fahimi, H.D., Mannaerts, G.P., van Veldhoven, P.P.: Molecular characterization of the human peroxisomal branched-chain acyl-CoA oxidase: cDNA cloning, chromosomal assignment, tissue distribution, and evidence for the absence of the protein in zellweger syndrome 93(24), 13748–13753. https://doi.org/10.1073/pnas.93.24.13748

[39] Russell, L., Garrett-Sinha, L.A.: Transcription factor ets-1 in cytokine and chemokine gene regulation 51(3), 217–226. https://doi.org/10.1016/j.cyto.2010.03.006. Accessed 2023-01-03

[40] Caron, C., Pivot-Pajot, C., van Grunsven, L.A., Col, E., Lestrat, C., Rousseaux, S., Khochbin, S.: Cdyl: a new transcriptional co-repressor 4(9), 877–882. https://doi.org/10.1038/sj.embor.embor917

[41] Dobin, A., Davis, C.A., Schlesinger, F., Drenkow, J., Zaleski, C., Jha, S., Batut, P., Chaisson, M., Gingeras, T.R.: STAR: ultrafast universal RNA-seq aligner 29(1), 15–21. https://doi.org/10.1093/bioinformatics/ bts635

[42] Bray, N.L., Pimentel, H., Melsted, P., Pachter, L.: Near-optimal probabilistic RNA-seq quantification 34(5), 525–527. https://doi.org/10.1038/nbt.3519. Number: 5 Publisher: Nature Publishing Group. Accessed 2022-12-10

[43] Quinlan, A.R., Hall, I.M.: BEDTools: a flexible suite of utilities for comparing genomic features 26(6), 841–842. https://doi.org/10.1093/bioinformatics/btq033. Accessed 2022-12-10

[44] Pezoa, F., Reutter, J.L., Suarez, F., Ugarte, M., Vrgoč, D.: Foundations of JSON schema. In: Proceedings of the 25th International Conference on World Wide Web, pp. 263–273. International World Wide Web Conferences Steering Committee. https://doi.org/10.1145/2872427.2883029. https://dl.acm.org/doi/10.1145/2872427.2883029 Accessed 2023-04-25

[45] Li, H.: Minimap2: pairwise alignment for nucleotide sequences 34(18), 3094–3100. https://doi.org/10.1093/bioinformatics/bty191. Accessed 2023-04-25

[46] Kreitzer, F.R., Salomonis, N., Sheehan, A., Huang, M., Park, J.S., Spindler, M.J., Lizarraga, P., Weiss, W.A., So, P.-L., Conklin, B.R.: A robust method to derive functional neural crest cells from human pluripotent stem cells 2(2), 119–131. Accessed 2022-11-09

[47] Herrmann, C.J., Schmidt, R., Kanitz, A., Artimo, P., Gruber, A.J., Zavolan, M.: PolyASite 2.0: a consolidated atlas of polyadenylation sites from 3 end sequencing 48, 174–179. https://doi.org/10.1093/nar/gkz918. Accessed 2023-02-08

[48] Klein, E., Ben-Bassat, H., Neumann, H., Ralph, P., Zeuthen, J., Polliack, A., Vánky, F.: Properties of the k562 cell line, derived from a patient with chronic myeloid leukemia 18(4), 421–431. https://doi.org/10.1002/ijc.2910180405

