## Supplementary methods for "SQANTI3: curation of long-read transcriptomes for accurate identification of known and novel isoforms"

#### SQANTI3 QC

##### STAR

If matching short-reads are provided to SQANTI3, it will run internally STAR and Kallisto to generate orthogonal information, which eventually serves to characterize the transcriptome and validate (or reject) transcript models during the curation process.

The exact commands used are the following:

```
STAR --runThreadN <num cpus> \
  --runMode genomeGenerate \
  --genomeDir <index_dir> \
  --genomeFastaFiles <fasta_genome> \
  --outTmpDir <index_dir_tmp>

STAR --runThreadN  <num cpus> --genomeDir <index_dir> \
    --readFilesIn <read1> <read2> \
    --outFileNamePrefix <sample_prefix> \
    --alignSJoverhangMin  8 \
    --alignSJDBoverhangMin 1 --outFilterType BySJout \
    --outSAMunmapped Within \
    --outFilterMultimapNmax  20 \
    --outFilterMismatchNoverLmax 0.04 \
    --outFilterMismatchNmax 999 --alignIntronMin 20 \
    --alignIntronMax 1000000 --alignMatesGapMax 1000000 \
    --sjdbScore 1 --genomeLoad NoSharedMemory \
    --outSAMtype BAM SortedByCoordinate --twopassMode Basic
```

##### Kallisto

We also implemented Kallisto's quantification inside SQANTI3 QC pipeline, so if pair-end data is input, Kallsito will create an index with the sequences of the transcript models being evaluated and quantify them using short reads. The exact commands are these ones:

```
kallisto index -i <kallisto_index> \
<corrected_fasta>  --make-unique

kallisto quant -i <kallisto_index> \
    -b 100 -t <num cpus> <read1> <read2>
```

The process is fully automated for pair-end data, but in some cases, like the K562 dataset where only single-end data is available, Kallisto (or any other quantification tool) can be run previously and provide its output to SQANTI3 via "--expression" parameter. In the K562 case, as the length of the fragments cannot be estimated with single-end reads, the same command described previously was run with some extra parameters based on the metadata available in the dataset ENCODE site:

```
kallisto quant -i <kallisto_index> \
    -b 100 -t <num cpus> --single \
```

```
    --rf-stranded -l 240 -s 30 <read1>
```

### Running IsoSeq3 from subreads to transcriptome

#### CCS

```
Subreads=”subreads/”$sample".bam"
ccs=$sample".ccs.bam"

ccs $subreads $ccs --min-rq 0.9
```

#### lima and refine

```
names=”cDNA_subreads_names.txt"
sample="WTC11.cDNA"
primers="primer.fasta"
fofn=$sample".flnc.fofn"
while read -r line
do
ccs="subreads/"$line".ccs.bam"
fl=$line".fl.bam"
echo "Running lima..."
date
lima $ccs $primers $fl --isoseq --peek-guess -j8

echo "Finished lima..."
date
fl_primers=$line".fl.Clontech_5p--Clontech_3p.bam"
flnc="IsoSeq3_output/"$line".flnc.bam"
echo "Runing refine..."
date
isoseq3 refine $fl_primers $primers $flnc --require-polya -j 8 --
verbose

echo "Finished refine..."
date
echo $flnc >> $fofn
done < $names
```

#### IsoSeq3 cluster and collapse

```
clustered=$sample".clustered.bam"
isoseq3 cluster $fofn $clustered --verbose --use-qvs -j 8

refGenome="lrgasp_grch38_sirvs.fasta"
hq_isoforms=$sample".clustered.hq.fasta"
hq_sam=$sample".clustered.hq.sam"
hq_sorted_sam=$sample".clustered.hq.sorted.sam"

minimap2 -ax splice -t 8 -uf --secondary=no -C5 $refGenome
$hq_isoforms > $hq_sam

sort -k 3,3 -k 4,4n $hq_sam > $hq_sorted_sam

collapse_isoforms_by_sam.py --input $hq_isoforms -s $hq_sorted_sam --
dun-merge-5-shorter -o $sample

get_abundance_post_collapse.py $sample".collapsed"
$sample".clustered.cluster_report.csv"
```

### SQANTI3 QC WTC11

#### Reference genome and annotation

```
refGenome="lrgasp_grch38_sirvs.fasta"
refGTF="lrgasp_gencode_v39_annotation_sirvs.human.gtf"
```

#### Short-read support

In order to speed up the data analysis and make it more consistent, we only ran once the mapping step with STAR using the command lines described above. Then, we took the resulting SJ.out.tab and BAM files to the rest of the analysis and input into SQANTI3 through the -c and –SR_bam arguments.

#### polyA motif list

This is the list of polyA motifs used as input via --polyA_motif_list :

```
aataaa
attaaa
agtaaa
tataaa
cataaa
gataaa
aatata
aataca
aataga
aaaaag
actaaa
aagaaa
aatgaa
tttaaa
aaaaca
ggggct
```

#### SQANTI3 QC for LI

```
python sqanti3_qc.py $sample $refGTF $refGenome \
       --fl_count $fl \
       -c $cov --short_reads $SR \
       --SR_bam $SR_bam \
       -o WTC11_cDNA.LI -d $out_dir -t8 \
       --report both -t 8
```

#### SQANTI3 QC for HIR & HIS

```
refTSS_peaks=human.refTSS_v3.1.hg38.bed # or WTC11_CAGE.filtered.bed
polyAsite=polyASite_atlas.2.0.GRCh38.96.bed
python sqanti3_qc.py $sample $refGTF $refGenome \
       --CAGE_peak $refTSS_peaks --fl_count $fl \
        --polyA_motif_list $polyA_motifs \
        -c $cov --short_reads $SR \
        --polyA_peak $polyA_sites --SR_bam $SR_bam \
        -o WTC11_cDNA.HIR -d $out_dir -t8 \
        --report both -t 8
```

#### SQANTI3 QC for K562

```
cage_peaks=CAGE_data.ENCFF981XPE.bed
polyA_sites=polyASite_atlas.2.0.GRCh38.96.bed

python sqanti3_qc.py $sample $refGTF $refGenome \
        --CAGE_peak $cage_peaks --fl_count $fl \
        --polyA_motif_list $polyA_motifs \
        --short_reads $SR --expression $kallisto_expression \
        --polyA_peak $polyA_sites_ref \
        -o K562_replicate1.SQ3 -d $out_dir -t8 \
        --report both
```

#### SQANTI3 QC for H1-endoderm

```
isoAnnot_gff4="Homo_sapiens_Gencode_v39.zip"
python sqanti3_qc.py $sample $refGTF $refGenome \
        --CAGE_peak $refTSS_peaks --fl_count $fl \
        --polyA_motif_list $polyA_motifs \
        --short_reads $SR \
        --polyA_peak $polyA_sites -e $expression_files \
        -o real_case.SQ3 -d $out_dir -t8 \
        --report both -t 8 --isoAnnotLite --gff3 $isoAnnot_gff3
```

#### SQANTI3 QC for reference

This is the command used to characterize the reference annotation regarding to the available data in each scenario. `cage_peaks_ref` and `polyA_sites` variables will change depending on the context.

```
python sqanti3_qc.py $refGTF $refGTF $refGenome \
        --CAGE_peak $cage_peaks_ref \
        --polyA_motif_list $polyA_motifs \
        --short_reads $SR --genename \
        --polyA_peak $polyA_sites \
        -o reference.SQ3 -d $out_dir -t8 \
        --report skip -t 8 --force_id_ignore --min_ref_len 0
```
